# Unique genomic sequences in a novel *Mycobacterium avium* subsp. *hominissuis* lineage enable fine scale transmission route tracing during pig movement

**DOI:** 10.1101/2022.04.04.487006

**Authors:** Tetsuya Komatsu, Kenji Ohya, Atsushi Ota, Yukiko Nishiuchi, Hirokazu Yano, Kayoko Matsuo, Justice Opare Odoi, Shota Suganuma, Kotaro Sawai, Akemi Hasebe, Tetsuo Asai, Tokuma Yanai, Hideto Fukushi, Takayuki Wada, Shiomi Yoshida, Toshihiro Ito, Kentaro Arikawa, Mikihiko Kawai, Manabu Ato, Anthony D. Baughn, Tomotada Iwamoto, Fumito Maruyama

**Affiliations:** Aichi Prefectural Tobu Livestock Hygiene Service Center, Toyohashi, Aichi, Japan; Faculty of Applied Biological Sciences, Gifu University, Gifu, Gifu, Japan; United Graduate School of Veterinary Sciences, Gifu University, Gifu, Gifu, Japan; Education and Research Center for Food Animal Health, Gifu University (GeFAH), Gifu, Gifu, Japan; Data Science Center, Division of Biological Science, Nara Institute of Science and Technology, Ikoma, Nara, Japan; Office of Academic Research and Industry-Government Collaboration, Hiroshima University, Higashi-Hiroshima, Hiroshima, Japan; Graduate School of Life Sciences, Tohoku University, Sendai, Miyagi, Japan; Kumamoto Prefectural Aso Public Health Center, Aso, Kumamoto, Japan; Division of Transboundary Animal Disease Research, National Institute of Animal Health, National Agriculture Research Organization, Tsukuba, Ibaraki, Japan; Toyama Prefectural Meat Inspection Center, Imizu, Toyama, Japan; Hiwa Natural History Museum, Shobara, Hiroshima, Japan; Graduate School of Human Life Science, Osaka City University, Osaka, Osaka, Japan; Clinical Research Center, National Hospital Organization Kinki-Chuo Chest Medical Center, Sakai, Osaka, Japan; Laboratory of Proteome Research, Proteome Research Center, National Institutes of Biomedical Innovation, Health and Nutrition, Ibaraki, Osaka, Japan; Department of Infectious Diseases, Kobe Institute of Health, Kobe, Hyogo, Japan; Graduate School of Human and Environmental Studies, Kyoto University, Kyoto, Kyoto, Japan; Department of Mycobacteriology, Leprosy Research Center, National Institute of Infectious Diseases, Higashimurayama, Tokyo, Japan; Department of Microbiology and Immunology, University of Minnesota Medical School, Minneapolis, Minnesota, USA; Project Research Center for Holobiome and Built Environment (CHOBE), Hiroshima University, Higashi-Hiroshima, Hiroshima, Japan; Scientific and Technological Bioresource Nucleus, Universidad de La Frontera, Temuco, Chile

**Author notes:** Corresponding author (FM). National Institute of Health Sciences, Kawasaki, Kanagawa, Japan (Kenji Ohya). Animal Research Institute, Council for Scientific and Industrial Research, Achimota, Accra, Ghana (Justice Opare Odoi). Central Research Institute for Feed and Livestock of Zen-noh, Tsukuba, Ibaraki, Japan (Shota Suganuma). Laboratory of Experimental Immunology, Department of Regeneration Science and Engineering, Institute for Life and Medical Sciences, Kyoto University, Kyoto, Japan (Toshihiro Ito). These authors contributed equally to this work (Tetsuya Komatsu, Kenji Ohya).

**Keywords:** MAH, VNTR, Draft genome sequences, pig, transmission route

## Abstract

*Mycobacterium avium* subsp. *hominissuis* (MAH) is one of the most prevalent mycobacteria causing non-tuberculous mycobacterial disease in humans and animals. Of note, MAH is a major cause of mycobacterial granulomatous mesenteric lymphadenitis outbreaks in pig populations. To determine the precise source of infection of MAH in a pig farm and to clarify the epidemiological relationship among pig, human and environmental MAH lineages, we collected 50 MAH isolates from pigs reared in Japan and determined draft genome sequences of 30 isolates. A variable number of tandem repeat analysis revealed that most pig MAH isolates in Japan were closely related to North American, European and Russian human isolates but not to those from East Asian human and their residential environments. Historical recombination analysis revealed that most pig isolates could be classified into SC2/4 and SC3, which contain MAH isolated from pig, European human and environmental isolates. Half of the isolates in SC2/4 had many recombination events with MAH lineages isolated from humans in East Asia. To our surprise, four isolates belonged to a new lineage (SC5) in the global MAH population. Members of SC5 had few footprints of inter-lineage recombination in the genome, and carried 80 unique genes, most of which were located on lineage specific-genomic islands. Using unique genetic features, we were able to trace the putative transmission route via their host pigs. Together, we clarify the possibility of species-specificity of MAH in addition to local adaptation. Our results highlight two transmission routes of MAH, one exposure on pig farms from the environment and the other via pig movement. Moreover, our study also warns that the evolution of MAH in pigs is influenced by MAH from patients and their residential environments, even if the MAH are genetically distinct.

**Highlights:**

- Variable number of tandem repeat analysis of *Mycobacterium avium* subsp. *hominissuis* (MAH) isolated from pigs (*n*=50) were conducted.
- Draft genome sequences of MAH (*n*=30) and genome analysis were conducted.
- Pig MAHs were genetically far from East Asian human isolates and close to those of Western countries.
- Novel MAH lineage which were transmitted farms by pig movement was found.
- Human MAH isolates influenced the evolution of pig isolates.

## Introduction

*Mycobacterium avium* complex (MAC) is the most common causative agent of non-tuberculous mycobacterial disease in humans in Asia and Europe (1, 2). The frequency of pulmonary MAC infection is increasing worldwide (3). *M. avium* subsp. *hominissuis* (MAH) is the most prevalent subspecies of MAC clinical isolates in Asia and Europe (1, 2).

MAH also causes lymphadenitis most often involving mesenteric lymph nodes in pigs (4, 5). Pigs infected with MAH usually have no clinical signs and the lesions are incidentally found during slaughter, bringing great financial losses in pig farm management in the event of an outbreak (6, 7).

In previous studies, IS*1245* or IS*1311* restriction fragment length polymorphism (RFLP) analysis revealed that environmental materials in pig farms such as sawdust (8) and peat (9) are important sources of MAH infection in pigs (10). The pig flow and the mother pigs are also important sources of MAH infection by RFLP analysis (11). However, discrimination power of RFLP analysis is inferior to a variable number of tandem repeat (VNTR) analysis (12).

Recently, several epidemiological studies using VNTR analysis of MAH have been described from around the world (4, 12, 13, 14, 15, 16, 17, 18, 19). VNTR analysis shows epidemiological relationships of MAH strains between humans and pigs (4, 16, 17, 18, 19). From these studies, it is suggested that MAH would have region-specific genetic diversity and distribution. In East Asian countries, human MAH strains isolated from patients with lung disease were identical to those from the residential environment, such as bathrooms or showerheads, and not identical to MAH isolates of pigs reared in the same region (4, 16). On the contrary, MAH strains of pigs in Japan were identical to those of humans from Europe (4, 13, 16,) and Japan (19). However, MAH strains of pigs in these studies were isolated from limited regions (Kinki, Kyushu and Okinawa area in Japan) (Supplementary Figure 1) (4, 13, 16, 19), in addition to the limitation due to use of only 8 loci for VNTR (19), which was thought to be less useful than 16 loci used in other analyses (12). To circumvent these shortcomings, MAH isolates from additional regions are needed to provide a more robust view of MAH distribution.

Phylogenetic and population structure analysis based on MAH genomes has recently revealed that MAH can be divided into six major clades, including MAHEastAsia1 (EA1), MAHEastAsia2 (EA2) and SC1-SC4, suggesting that MAH adaptation is geographically restricted around the world (20, 21). For example, EA1 and EA2 lineages were mainly distributed in humans in Japan and Korea although SC1-SC4 lineages were distributed in Europe and USA (20, 21). Although these studies utilized many MAH genomes, only two pig isolates were included. Therefore, by using comparative genome analysis of additional pig MAH isolates, it is possible to unravel the relationship between human, pig and environment MAH isolates, and decipher respective transmission routes.

In this study, we aimed to characterize the distribution of MAH lineages in farmed pigs in Japan and their genetic relationship with human and environmental MAH isolates for better understanding of its diversity and the way of infection. In addition, we collected detailed epidemiological information for the determination of transmission route of MAH for pigs.

## Materials and methods

### Bacterial isolation and identification

The approach used for isolation and identification of MAH is described in our previous study (22). 50 isolates used for VNTR analysis were isolated from lymph nodes, liver, feces, sawdust and farm soil (Supplementary Table 1). All of them were isolated from pigs or farm environments in Tokai area (Gifu, Aichi and Shiga Prefecture) and Hokuriku area (Toyama and Ishikawa Prefecture), whose geographical relationship is depicted in Supplementary Figure 1.

### Mycobacterial Interspersed Repetitive Unit and *Mycobacterium avium* Tandem Repeat VNTR

Primer sets for 15 loci of *Mycobacterium avium* Tandem Repeat (MATR) (MATR1-9, 11-16) and 7 loci selected from Mycobacterial Interspersed Repetitive Unit (MIRU) loci (32, 292, X3, 25, 7, 10, 47) were used to generate VNTR profiles, as previously reported (16). The data profiles of 50 MAH isolates in this study and the number of VNTR profiles were available in Supplementary Table 2 and 3. From all data, we selected 13 MATR (MATR1-8, 11-13, 15-16) and 7 MIRU profiles for minimum spanning tree (MST) analysis to compare with variable data of past reports (4, 13, 14, 16, 19, 23, 24) using Bionumerics version 7.0 (Applied Maths, Sint-Martens-Latem, Belgium). Clonal complex (CC) in MST of 13 loci was defined as a different group when there were two or more differences in the VNTR profile of adjacent nodes according to the past study with one modify, that a CC was composed when it had over five genotypes (13). As the discrimination ability of MST analysis based on MIRU-VNTR was too low, we were not able to divide the MAH isolates into several CCs and therefore did not show a corresponding index.

### Draft genome sequence

Draft genome sequences of 30 isolates were obtained according to our past study (22).

#### Genome phylogenetic tree and single nucleotide polymorphism analysis of *cinA* and *sugA* gene

Genome phylogenetic tree analysis based on 30 MAH isolates and 20 reference strains (21) were performed via the online tool CSI Phylogeny version 1.4 with default parameters (25) and IQ-TREE (26) by maximum likelihood method. *M. avium* subsp. *paratuberculosis* (MAP) was set as an outgroup. Obtained newick file was visualized by Molecular Evolutionary Genetics Analysis (MEGA) 7.0. Alignment file generated by CSI Phylogeny was used for phylogenetic analysis by neighbor joining method with p-distance method via MEGA 7.0. Topological evaluation of obtained newick file was conducted by IQ-TREE (26). All of the reference genome sequences were retrieved from National Center for Biotechnology Information (NCBI) genome database. The sequence of two epidemiological marker genes, *cinA* (product name: cytochrome P-450) and *sugA* (product name: trehalose ABC transporter), which are offered to be able to distinguish major six MAH lineages (21), were retrieved from draft genome sequences. These sequences and the reference strains were aligned and used for the phylogenetic tree analysis by maximum likelihood method via MEGA 7.0 and single nucleotide polymorphism (SNP) analysis.

#### Bayesian analysis of population structure and recombination analysis for determination of MAH lineage

MAH lineage was predicted by bayesian analysis of population structure (BAPS) using fastBAPS (27) and fastGEAR (28), which was able to infer the MAH lineages, considering genetic linkage (21, 27). Used genome sequences were the same as using for followed phylogenetic tree analysis except for MAP. Filtered polymorphic sites in vcf file, which contained the information of genetic variants against reference genome sequences, generated by CSI Phylogeny were used for fastBAPS (27). In addition, these sites were combined with intervening reference genome sequences TH135, and were organized in an alignment file (21) and then used as an input for fastGEAR (28). Lineages were then generated by hierarchical clustering using fastGEAR (28), which produced lineage classifications that were highly consistent with previous reports (20, 21).

#### Pan and core genome analysis

Draft genome sequences, which were the same as BAPS and recombination analysis, were re-annotated to obtain gff files by Prokka v1.14.5 (29). Genes present or absent only in four isolates (GM10, GM12, GM21, GM32), which we named SC5 as a new lineage below, were searched by Roary v3.11.2 (30). Basic Local Alignment Search Tool (BLAST) analysis was performed with the genes present only in SC5 and annotated the genes which had high coverage and similarity (over 95%) with genes of MAH in NCBI gene database. We also evaluated whether these genes were located on genomic islands and/or phage regions as described below.

#### Detection of mobile elements (Plasmid, phage and genomic island) in MAH genomes

Plasmids were identified from draft genome sequences by PlasmidFinder v 2.1 with the following settings (threshold for minimum % identity: 50%, minimum % coverage: 20%) (31). Prophage sequences were identified by PHASTER (PHAge Search Tool Enhanced Release) (32, 33). Genomic islands were predicted by IslandViewer version 4 (34).

#### Epidemiological information of farms

Epidemiological information of each farm about bedding materials, introduction and shipping of pigs was obtained via interviewing farm owners.

## Results

### Mycobacterial Interspersed Repetitive Unit and *Mycobacterium avium* Tandem Repeat VNTR

#### MIRU VNTR based on 7 loci

MAH isolates were classified into three groups (Supplementary Figure 2). Group 1 contained mainly the human isolates, although almost all pig isolates in Japan were classified in Group 2. Several pig isolates were classified in Group 1, however, no isolate was clustered with human isolates. The pig isolates formed a cluster with environmental isolates in Group 2. The eight isolates of including four SC5 isolates formed Group 3.

#### MATR VNTR based on 13 loci

Isolates were divided into six CCs (Figure 1). Most human isolates in Japan and Korea were classified into CC1. CC2 was composed of MAH isolates of USA, European, Russian and some Korean human and Japanese pig isolates. CC4 and CC5 were mainly composed of pig isolates from Japan, although CC3 contained human and pig isolates from Japan. The isolates same as group 3 in MST of 7 loci were classified into CC6 and were genetically far from any adjacent node.

**Figure 1.**
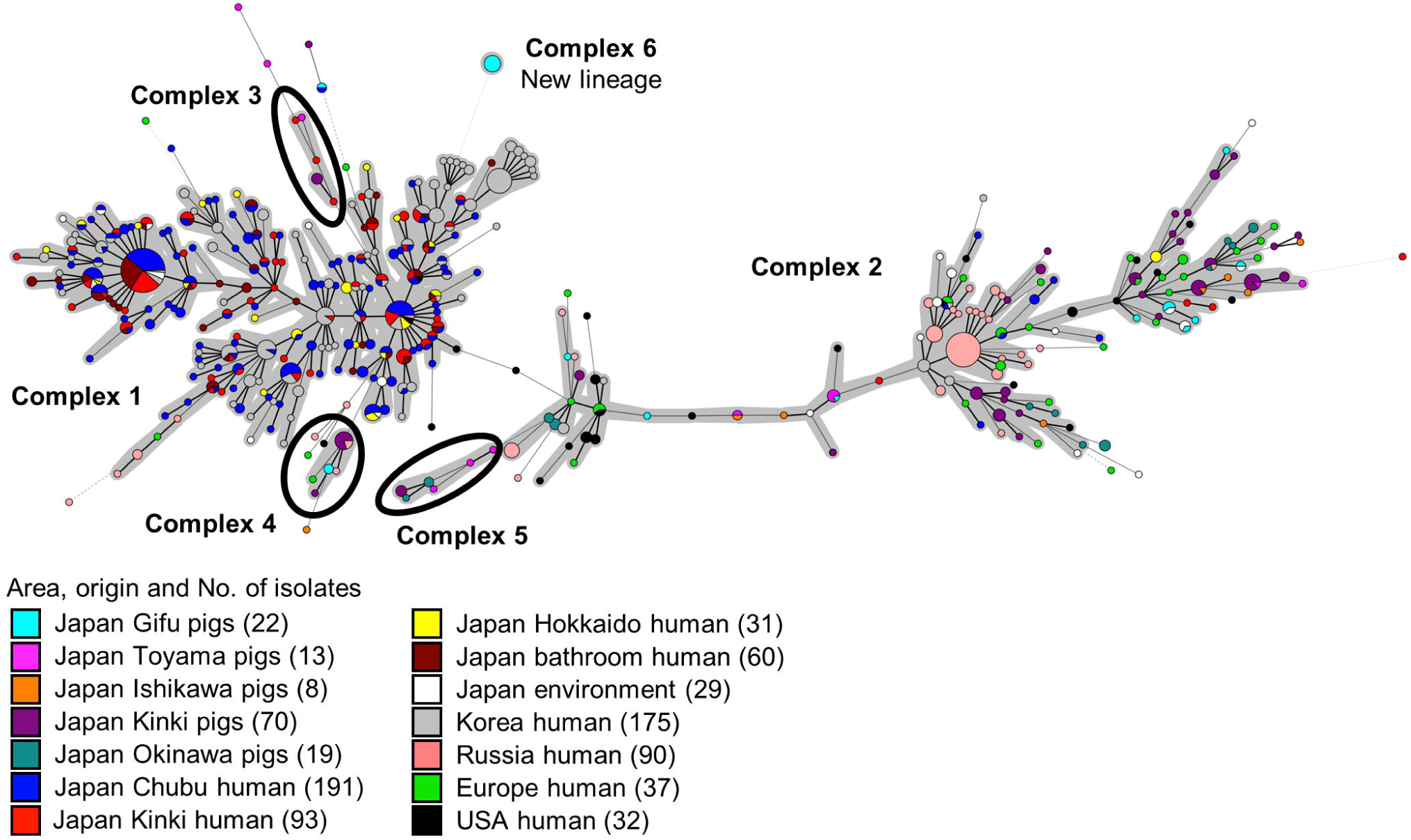
Minimum spanning tree based on 13 loci *Mycobacterium avium* Tandem Repeat VNTR genotyping of MAH isolates from various geographic regions and sources. Circles indicate different VNTR profiles. The size of each circle depended on the number of isolates sharing the same profiles. Clonal complexes were highlighted in gray. Each clonal complex consisted of over five isolates.

#### Phylogenetic tree analysis

As a result of topological evaluation of both phylogenetic trees, the phylogenetic tree of CSI phylogeny (25) was considered to be more appropriate than that of the IQ-TREE (26). 24 isolates were classified into 3 main lineages (SC2: 13, SC3: 1, SC4: 10), and 4 isolates were classified into a new lineage (SC5) (Figure 2). SC5 was the earliest branch from the MAP (Figure 2, Supplementary Figure 3 and 4). GM16 and OCU484 were unclassified and placed between EA2 and SC3 (Figure 2, Supplementary Figure 3), however, they were classified in EA2 by the IQ-TREE (Supplementary Figure 4).

**Figure 2.**
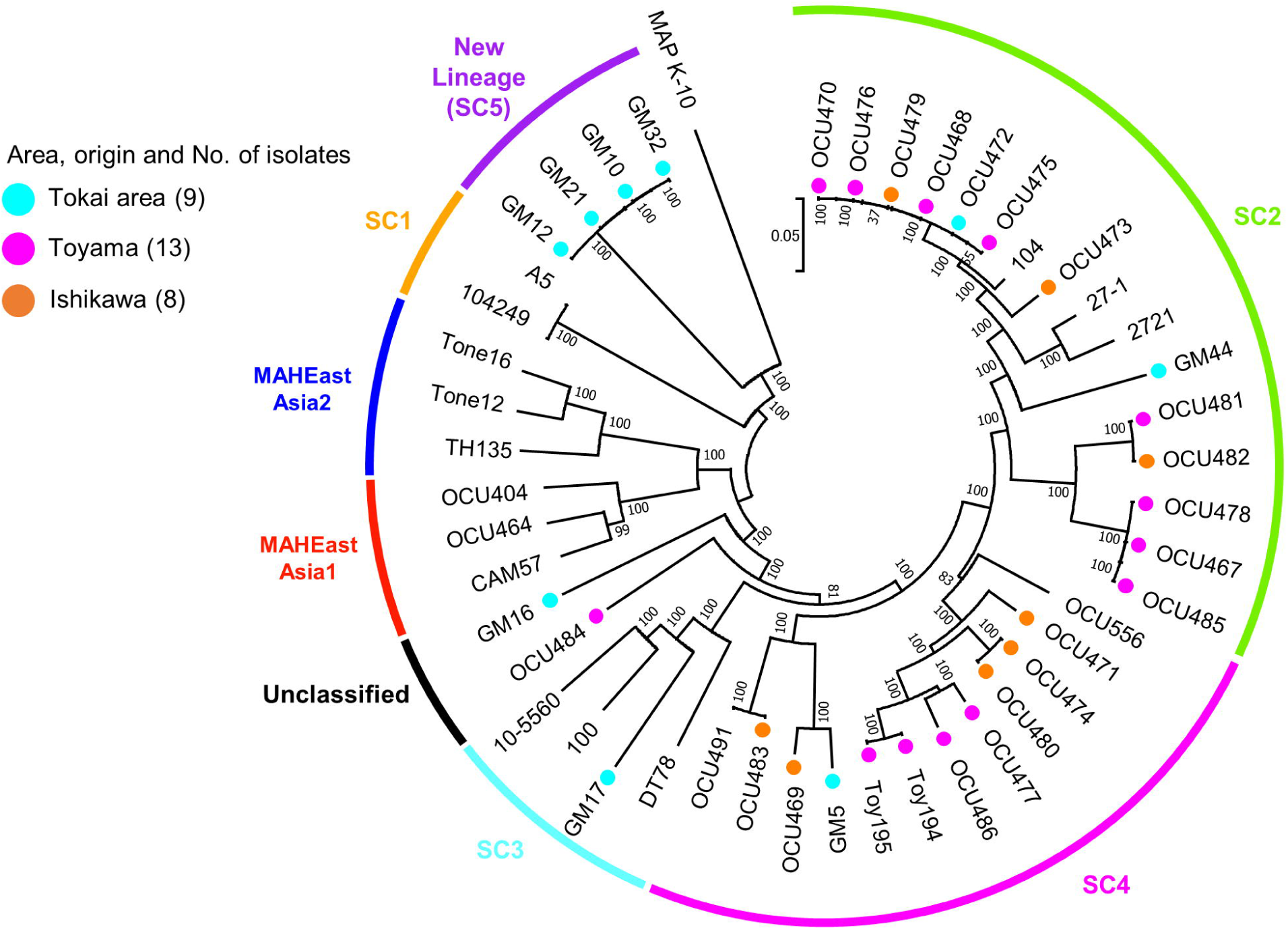
Phylogenetic tree based on draft genome sequences. Neighbor joining tree was generated by MEGA 7.0 using alignment file obtained from CSI phylogeny version 1.4. Bootstrap values were calculated by 1,000 replications. The scale bar indicates genetic distance of each strain. Lineage classification except for new lineage was based on Yano et al (21).

#### Phylogenetic analysis for *cinA* and *sugA* genes

SC5 shared a unique SNP pattern in both genes compared with the other lineage (Supplementary Figure 5 and 6). GM5 had a different SNP pattern from SC4 in *cinA* (Supplementary Figure 5). GM16 had the same SNP patterns as EA1 in *cinA*. GM16 and OCU484 had unique SNP patterns in *sugA*, respectively (Supplementary Figure 6). Interestingly, SNP in the *sugA* gene of SC2 were divided into two patterns, SC2a and SC2b. SNP patterns of SC2a were of a new type and had a characteristic SNP, C25T. The patterns of SC2b were the same as *M. avium* subsp *avium*.

#### Bayesian analysis of population structure and recombination analysis for lineage classification

The isolates were divided into 4 lineages, SC5, SC1, SC2 and large SC by fastBAPs (27) (Supplementary Figure 7). The occurrence of inter-lineage recombination events assessed by fastGEAR (28) is shown in Figure 3. The strains and isolates were divided into 5 lineages (SC5, SC1, EastAsia, SC2/4, and SC3). Each of the pig isolates was classified into three lineages (SC3: 1 isolate, SC2/4: 25, SC5: 4). SC5 had few recombination events with any other lineages. Similarly, some isolates of lineage SC2/4, which were classified into SC2a by SNP analysis, also had few occurrences of recombination among lineages.

**Figure 3.**
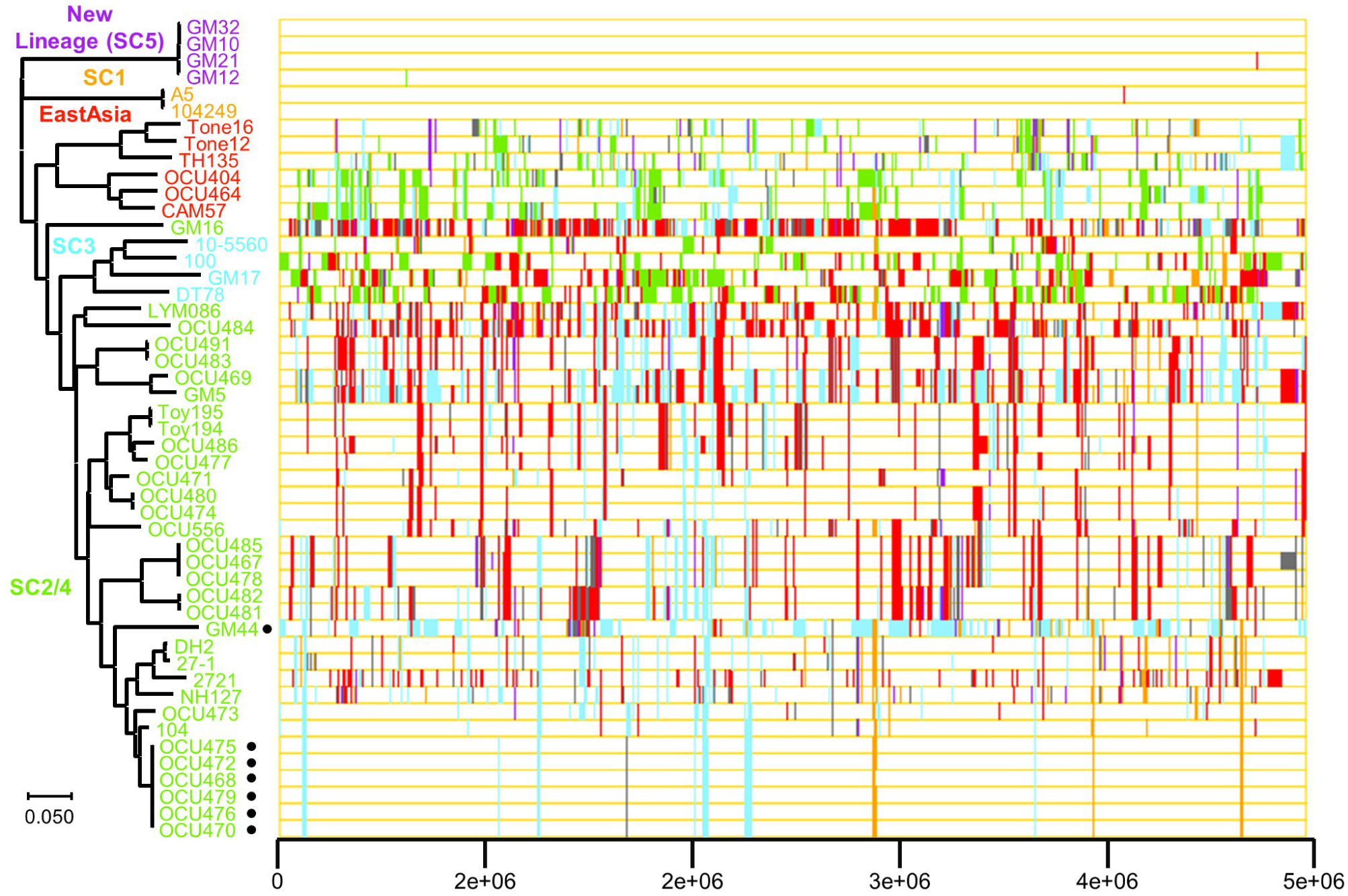
Determination of MAH lineages. Right: Visualization of recent horizontal acquisition events generated by fastGEAR. Color of chromosomal regions expresses donor lineage of the regions. Left: Neighbor joining tree was generated by via MEGA 7.0 using alignment file obtained from CSI phylogeny version 1.4.

#### Pan and core genome analysis

Core and accessory gene content were 2,314 and 12,115, respectively. Six core genes were absent only in the SC5 and 97 genes were characteristic in this lineage (Supplementary Table 4). From BLAST analysis, 80 genes of the SC5 had less than 95% coverage and similarity with the other MAH isolates. Most of them were clustered and were located on genomic islands, which were not previously reported (35).

#### Detection of mobile elements (Plasmid, phage and genomic island) in MAH genome

No plasmids were detected in any isolates. Intact and questionable phage regions were also not detected, although incomplete sequences of phage were detected in all isolates. Genomic islands were detected in all isolates, however, there was diversity in the number of genomic islands (Table 1).

**Table 1.** Predicted genomic islands and a number of genes on the genomic island.

| Isolates | Number of genomic islands | Number of genes in genomic island |
| --- | --- | --- |
| GM5 | 43 | 464 |
| GM10 | 37 | 454 |
| GM12 | 35 | 485 |
| GM16 | 51 | 565 |
| GM17 | 43 | 391 |
| GM21 | 36 | 434 |
| GM32 | 34 | 424 |
| GM44 | 45 | 638 |
| OCU467 | 7 | 89 |
| OCU468 | 9 | 253 |
| OCU469 | 28 | 321 |
| OCU470 | 7 | 217 |
| OCU471 | 35 | 311 |
| OCU472 | 8 | 180 |
| OCU473 | 38 | 489 |
| OCU474 | 41 | 459 |
| OCU475 | 9 | 164 |
| OCU476 | 10 | 179 |
| OCU477 | 35 | 376 |
| OCU478 | 8 | 122 |
| OCU479 | 11 | 295 |
| OCU480 | 37 | 462 |
| OCU481 | 8 | 111 |
| OCU482 | 7 | 93 |
| OCU483 | 11 | 155 |
| OCU484 | 32 | 451 |
| OCU485 | 8 | 112 |
| OCU486 | 40 | 379 |
| Toy194 | 44 | 686 |
| Toy195 | 50 | 624 |

#### Farm relationship where the novel lineage MAH was isolated

The isolates of SC5 were isolated from pigs reared in three farms located in Gifu, Shiga and Aichi Prefecture (Supplementary Table 1). From interviewing farm owners, these farms were related epidemiologically (Figure 4). Aichi B and Gifu A farms had introduced boars and gilts from Aichi Y farm. Shiga A farm had introduced fattening pigs from Aichi X farm, whose gilts were introduced from Aichi Y farm. There was nothing in common about using animal bedding materials, such as sawdust, among farms. Further, the other farms including in the Hokuriku area had not introduced pigs from the Tokai area.

**Figure 4.**
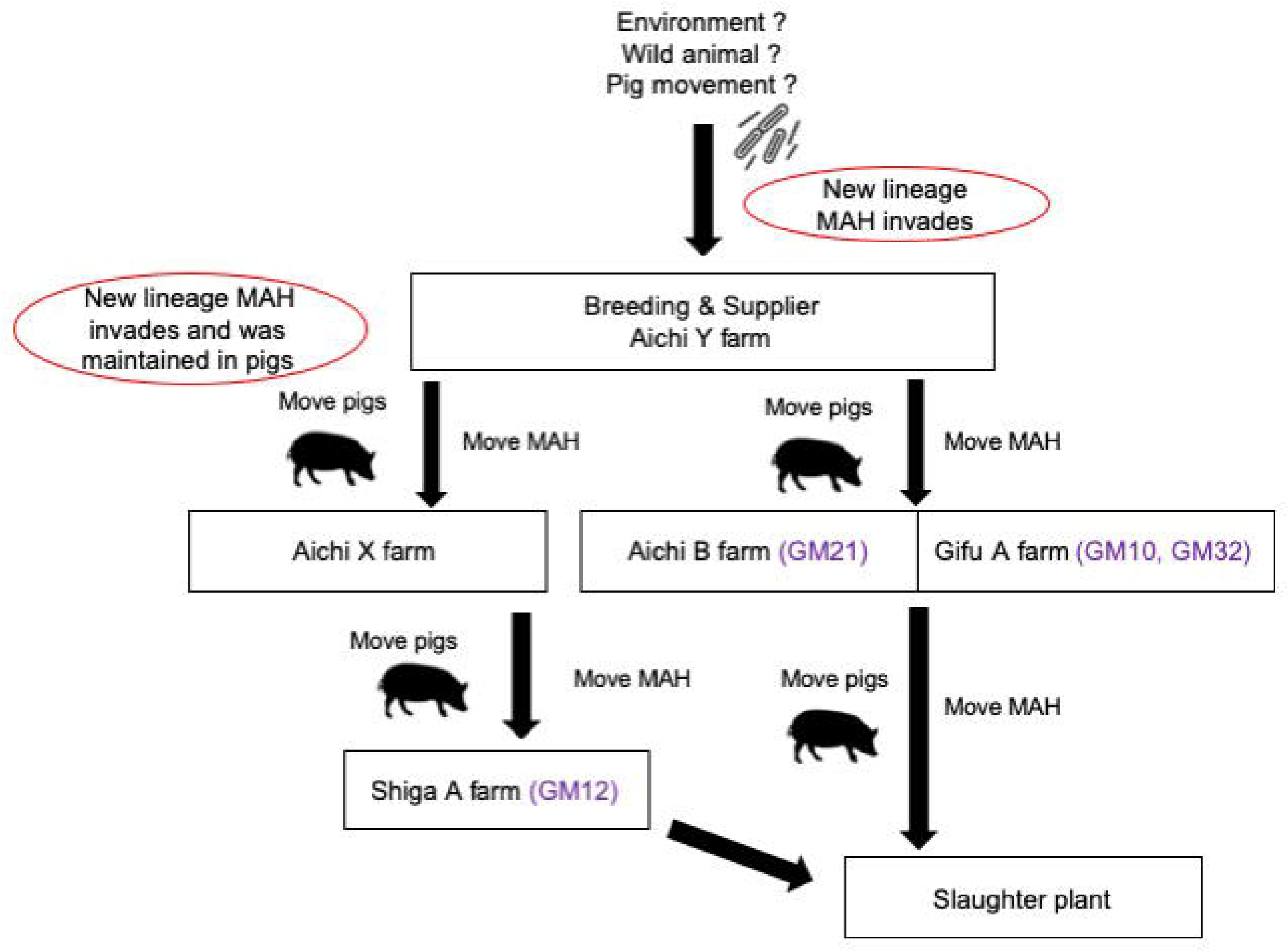
Relationship among farms where new lineage MAH was isolated. Aichi Y farm bred and supplied pigs to Aichi X, Aichi B and Gifu A farm, where these farms had the novel lineage (GM21, GM10 and GM32). Aichi X farm whose gilts were introduced from Aichi Y farm supplied pigs to Shiga A farm where GM12 was isolated. Finally, pigs of Gifu A, Aichi B and Shiga A farm were slaughtered in the same slaughtered plant (Gifu meat inspection center).

## Discussion

In the present study, we revealed three critical points, the first was the genetic relationship among MAH isolates of pigs, humans and the environment in Japan, the second was the identification of a new MAH lineage and its genomic and epidemiological features, and the third was putative transmission routes of pig MAH.

### Genetic relationship of pig isolates in Japan

From VNTR analysis using both 7 and 13 loci, most pig isolates were grouped with those from pigs and the environment in Japan and humans in Europe, Russia and USA, supporting previous studies (4, 13, 14, 16). Nevertheless, pig MAH isolates from various regional origins are closely genetically related, except for the new lineage. This observation indicates that genetically close MAHs are widely distributed among Japanese pigs.

In addition, by phylogenetic tree analysis based on draft genome sequences, 28 isolates were classified into SC2, SC3, SC4, SC5, and two isolates were unclassified. These results are consistent with previous reports (20, 21), but it is worth noting that SC3 was isolated from pigs in Japan for the first time (Supplementary Table 5). Recombination analysis clarified two previously unclassified isolates as belonging to lineage SC2/4. These isolates had many recombination events with ‘EastAsia’ and were classified into EA1 and SC4 by SNP analysis of the *cinA* gene, respectively. Frequent recombination events and multiple SNP in recombination-cold regions may contribute to the result of these isolates being unclassified in genome-based phylogenetic tree analysis. These isolates can be reclassified when several isolates with similar genetic features were found.

On the other hand, SNP analysis reveals there is diversity in pig isolates in Japan at the level of a single gene. SC2/4 isolates were divided into 5 patterns by phylogenetic analysis of *sugA*. They had been isolated from multiple farms across the prefectures. Focusing on SC2a, the isolates had a unique SNP pattern in their core gene and had few recombination events with the other lineages. This indicates that SC2a has evolved through accumulation of genetic variations by microevolution to facilitate to adapt itself to living pigs, rather than has evolved through recombination events in the environment.

In lineage SC4, most isolates had the same SNP pattern as the reference strain and had relatively frequent recombination events among lineages, consistent with a past study (21). SC4 was isolated from human and the environment in Europe, USA and Russia (21) (Supplementary Table 5) and Japanese pig isolates are genetically close to those of environmental isolates in these areas (13). According to a recent annual report from the Animal Quarantine Service in Japan, most live pigs that are imported to Japan originate from Canada (43.7%), USA (33.4%) and Denmark (12.9%) in 2019 (https://www.maff.go.jp/aqs/tokei/toukeinen.html). These observations suggest that SC4 may be transmitted from overseas countries by a pig movement and has evolved through recombination events with the ‘EastAsia’ lineage that is released from humans or the residential environments in Japan.

### Genomic and epidemiological features of the novel lineage

Four isolates of SC5 had the same characteristic SNP patterns in the *cinA* and *sugA* genes, VNTR profiles and were present in a very unique place in MST trees. These results including genome based phylogenetic tree, BAPS and recombination analysis indicated that this cluster was a novel lineage. SC5 isolates had 80 unique genes in their genomes, most of which were clustered on genomic islands. In addition, these genes contained mobile element associated genes, such as IS*481* family transposase IS*3514* and putative prophage phiRv2 integrase. SC5 had few recombination events with the other lineages and had characteristic SNPs in their core genes. These results suggest that SC5 can be partially characterized by mobile elements rather than recombination events (20, 21) and that they may adapt to the environment in living pigs via accumulation of genetic variations by microevolution. These unique genes may serve as the understanding of MAH survival strategies in living pigs.

It was unknown why SC5 was isolated only from the Tokai area in Japan. In general, the *Mycobacterium* genus is mainly present in the environment, such as in water sources, soil and dust (36, 37), however, SC5 is genetically distinct from the other MAH isolates including environmental, human and pig isolates from the world. MAH is isolated from wild boars in Europe (38), however, MAH is not isolated from wild animals in the same area of this study (39). Based on these observations, it is unlikely that SC5 was introduced from other areas or the environment. The epidemiological information suggests that SC5 is spreading via pig movement. This hypothesis is also supported by the observation that there were few inter-lineage recombination events in SC5, because it is thought that acquisition of genetic variation through recombination (20) or plasmid transfer (40) occurs more frequently in the environment than in living hosts. Our results also indicate that SC5 has species-specificity for pigs, suggesting the possibility of adaptation of MAH for specific environments including within living hosts and that there are likely additional lineages that have yet to be identified.

### Putative transmission route of pig MAH

We clarified two transmission route of pig MAH by recombination and SNP analysis, from environment and via pig movement (Figure 5). SC2a and SC5 would be transmitted farms via pig movement because they had few recombination events and characteristic SNP, although SC4 would be invaded into farms from the environment. Our study highlights that the elimination of subclinically infected pigs with MAH and disinfection of equipment brought from outside the pigsty could be effective tools against MAH invasion.

**Figure 5.**
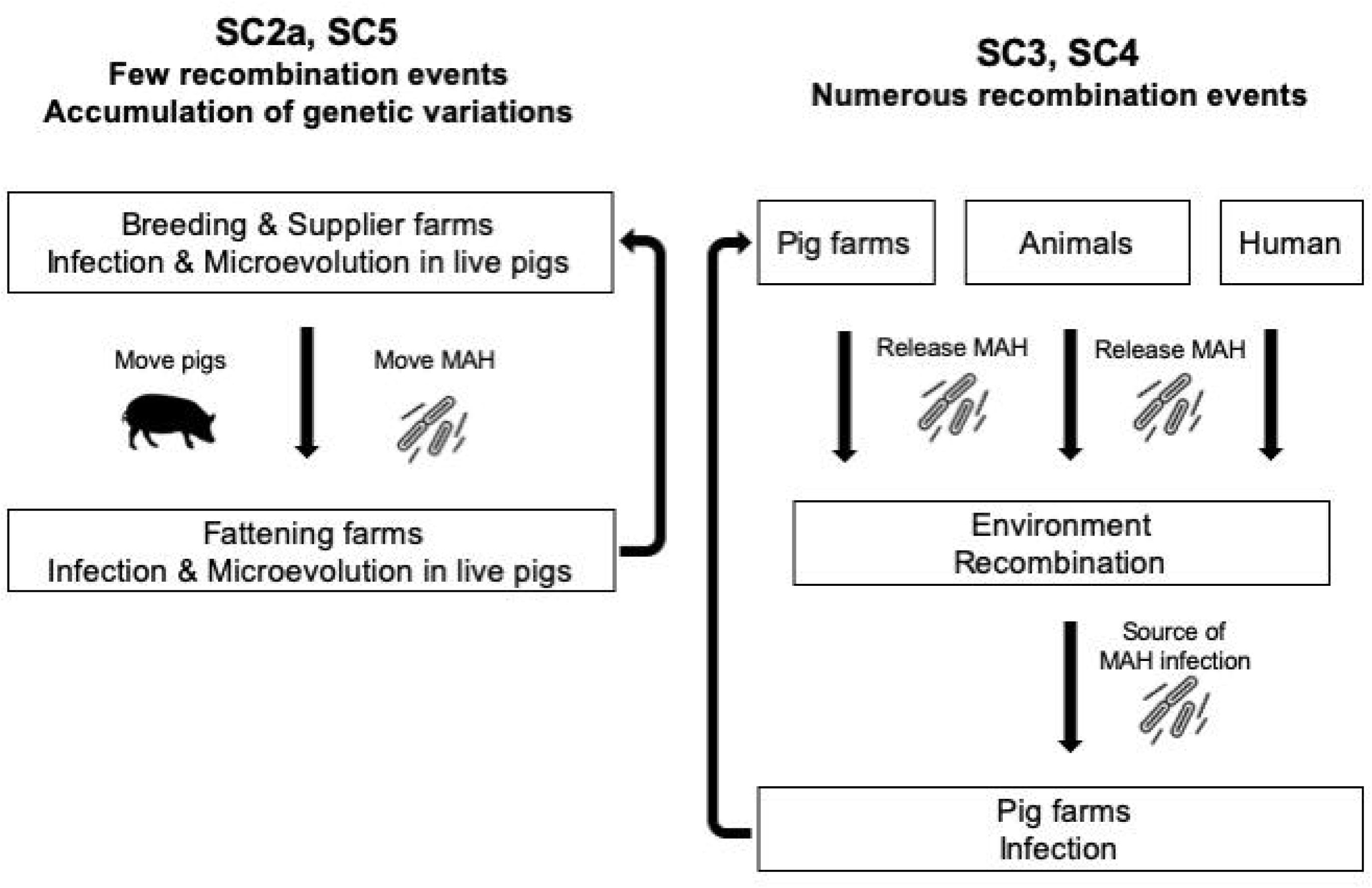
The differences in the way of MAH infection in pigs by lineages. SC2 and SC5 exist in live pigs and would evolve with the accumulation of genetic variations in their genes by microevolution. They move from farms to farms via live pigs. SC3 and SC4 evolve through numerous recombination events in the environment with the other lineages released from various hosts. MAH that have completed recombination would be supplied to pig farms from the environment.

## Conclusion

We have clarified the relationship among MAH isolates of pigs and humans in Japan and the world using VNTR and genome analysis. Our study highlights that there are two MAH transmission routes in pigs, via pig movement and from the environment. We have also identified that SC2a and SC5 can have evolved in a pig-specific manner. Finally, our study emphasizes that MAH in humans and the residential environments has become a source of transmission to the environment and is important for the evolution of pig MAH.

## Data availability

The draft genome sequence data used in this study have been submitted to the NCBI genome database (https://www.ncbi.nlm.nih.gov/genome/) and all of accession number was available in our past study (22).

## CRediT authorship contribution statement

Tetsuya Komatsu: Methodology, Software, Validation, Formal analysis, Investigation, Data Curation, Writing-Original Draft. Kenji Ohya: Conceptualization, Methodology, Validation, Resource, Data Curation, Writing-Review & Editing, Supervision, Project administration, Funding acquisition. Atsushi Ota: Software, Yukiko Nishiuchi: Conceptualization, Methodology, Resource, Writing-Review & Editing. Hirokazu Yano: Software, Writing-Review & Editing, Kayoko Matsuo: Resource, Data Curation, Investigation. Justice Opare Odoi: Methodology, Software, Formal analysis, Investigation, Writing-Review & Editing. Shota Suganuma: Methodology, Formal analysis, Investigation. Kotaro Sawai: Methodology, Formal analysis, Investigation, Writing-Review & Editing. Akemi Hasebe: Resource, Data Curation, Investigation. Tetsuo Asai: Writing-Review & Editing, Supervision. Tokuma Yanai: Writing-Review & Editing, Supervision. Hideto Fukushi: Writing-Review & Editing, Supervision. Takayuki Wada: Conceptualization, Methodology, Supervision. Shiomi Yoshida: Conceptualization, Methodology, Supervision. Toshihiro Ito: Software, Data Curation. Kentaro Arikawa: Conceptualization, Methodology. Mikihiko Kawai: Software, Data Curation, Writing-Review & Editing. Manabu Ato: Writing-Review & Editing, Supervision, Project administration. Anthony D. Baughn: Writing-Review & Editing, Supervision, Project administration. Tomotada Iwamoto: Conceptualization, Methodology, Writing-Review & Editing, Supervision, Project administration, Fumito Maruyama: Conceptualization, Methodology, Validation, Data Curation, Writing-Review & Editing, Supervision, Project administration, Funding acquisition.

## Acknowledgments

We thank the member of Gifu Prefectural Chuo Livestock Hygiene Service Center for sampling. We thank the Data Integration and Analysis Facility at the National Institute for Basic Biology for providing some of the computational resources.

## Supporting Information Legends

**Supplementary Table 1. Information of MAH isolates in this study.**

**Supplementary Table 2. VNTR profiles of 50 MAH isolates in this study.**

**Supplementary Table 3. Information of MAH strains or isolates used for VNTR analysis in this study.**

**Supplementary Table 4. Uniquely present or absent genes in new lineage (SC5).**

**Supplementary Table 5. Geographic and host features of MAH lineages.**

**Supplementary Figure 1. Location of each regions and prefectures in Japan.** Pigs from Tokai area (southern Gifu, Shiga and Aichi Prefecture) are slaughtered in Gifu meat inspection center. Pigs from Hokuriku area (northern Gifu, Toyama and Ishikawa Prefecture) are slaughtered in Toyama meat inspection center.

**Supplementary Figure 2. Minimum spanning tree (MST) based on 7 loci Mycobacterial Interspersed Repetitive Unit VNTR genotyping of MAH isolates in Japan.** Circles indicate different VNTR profiles. The size of each circle depended on the number of isolates sharing the same profiles. New lineages indicated by an arrow were clustered and consisted of isolates GM10, GM11, GM12, GM21, GM23, GM24, GM25, GM32.

**Supplementary Figure 3. Phylogenetic tree based on draft genome sequences generated by CSIphylogeny.** Maximum likelihood tree was generated by CSI phylogeny version 1.4 and was visualized via MEGA 7.0. The area and Prefecture of isolates in this study were indicated by the color dots. Lineage classification except for new lineage was based on Yano et al., 2019. BMC Genomics.

**Supplementary Figure 4. Phylogenetic tree based on draft genome sequences generated by IQ-TREE.** Maximum likelihood tree was generated by IQ-TREE and visualized via MEGA 7.0. The Area and Prefecture of isolates in this study were indicated by the color dots. Lineage classification except for new lineage was based on Yano et al., 2019. BMC Genomics.

**Supplementary Figure 5. Allelic variants in *cinA* gene of pig MAH.** A) Maximum likelihood tree generated by MEGA 7.0. Bootstrap values were calculated by 1,000 replications. The scale bar indicates genetic distance of each strain. Lineage classification except for new lineage was based on Yano et al (25). B) SNP cites in pig MAH. New lineage shared the same SNP patterns, which were highlighted by gray. Some variations were detected in SC4. Reference SNP patterns were referred from Yano et al (25).

**Supplementary Figure 6. Allelic variants in *sugA* gene of pig MAH.** A) Maximum likelihood tree generated by MEGA 7.0. Bootstrap values were calculated by 1,000 replications. The scale bar indicates genetic distance of each strain. Lineage classification except for new lineage was based on Yano et al (25). B) SNP cites in pig MAH. New lineage shared the same SNP patterns, which were highlighted by gray. Two type of SNP patterns were detected in SC2. Reference SNP patterns were referred from Yano et al (25).

**Supplementary Figure 7. Lineage classification by fastBAPS.** Lineage classification was predicted by fastBAPS (Tonkin-Hil et al., 2019 Nucleic Acids Res). Bootstrap values were calculated by 1,000 replications. The scale bar indicates genetic distance of each strain. Lineage classification in the right side except for new lineage was based on Yano et al., 2019. BMC Genomics. Neighbor joining tree was generated by via MEGA 7.0 using alignment file obtained from CSI phylogeny version 1.4.

## References

1. D.R. Prevots, T.K. Marras, Epidemiology of human pulmonary infection with nontuberculous mycobacteria: a review. Clin. Chest. Med. 36 (2015) 13–34.

2. H. Namkoong, A. Kurashima, K. Morimoto, Y. Hoshino, N. Hasegawa, M. Ato, S. Mitarai, Epidemiology of pulmonary nontuberculous mycobacterial disease, Japan. Emerg. Infect. Dis. 22 (2016) 1116–1117.

3. C.L. Daley, *Mycobacterium avium* Complex Disease. Microbiol. Spectr. 5 (2017).

4. T. Adachi, K. Ichikawa, T. Inagaki, M. Moriyama, T. Nakagawa, K. Ogawa, Y. Hasegawa, T. Yagi, Molecular typing and genetic characterization of *Mycobacterium avium* subsp. *hominissuis* isolates from humans and swine in Japan. J. Med. Microbiol. 65 (2016) 1289–1295.

5. A. Agdestein, T.B. Johansen, Ø. Kolbjørnsen, A. Jørgensen, B. Djønne, I. Olsen, A comparative study of *Mycobacterium avium* subsp. *avium* and *Mycobacterium avium* subsp. *hominissuis* in experimentally infected pigs. BMC. Vet. Res. 8 (2012) 11.

6. J. Álvarez, E. Castellanos, B. Romero, A. Aranaz, J. Bezos, S. Rodríguez, A. Mateos, L. Domínguez, L. de Juan. Epidemiological investigation of a *Mycobacterium avium* subsp. *hominissuis* outbreak in swine. Epidemiol. Infect. 139 (2011) 143–148. doi: 10.1017/S0950268810001779.

7. N. Hulinova Stromerova, M. Faldyna. Mycobacterium avium complex infection in pigs: A review. Comp. Immunol. Microbiol. Infect. Dis. 57 (2018) 62–68. doi: 10.1016/j.cimid.2018.06.005.

8. L. Matlova, L. Dvorska, K. Palecek, L. Maurenc, M. Bartos, I. Pavlik, Impact of sawdust and wood shavings in bedding on pig tuberculous lesions in lymph nodes, and IS1245 RFLP analysis of *Mycobacterium avium* subsp. *hominissuis* of serotypes 6 and 8 isolated from pigs and environment. Vet. Microbiol. 102 (2004) 227–236.

9. A. Agdestein, T.B. Johansen, V. Polaček, B. Lium, G. Holstad, D. Vidanović, S. Aleksić-Kovačević, A. Jørgensen, J. Žultauskas, S.F. Nilsen, B. Djønne B, Investigation of an outbreak of mycobacteriosis in pigs. BMC. Vet. Res. 7 (2011) 63.

10. I.A. Gardner, D.W Hird. Environmental source of mycobacteriosis in a California swine herd. Can. J. Vet. Res. 53 (1989) 33–37.

11. K. Hibiya, Y. Kazumi, Y. Nishiuchi, I. Sugawara, K. Miyagi, Y. Oda, E. Oda, J. Fujita. Descriptive analysis of the prevalence and the molecular epidemiology of *Mycobacterium avium* complex-infected pigs that were slaughtered on the main island of Okinawa. Comp. Immunol. Microbiol. Infect. Dis. 33 (2010) 401–421.

12. T. Inagaki, K. Nishimori, T. Yagi, K. Ichikawa, M. Moriyama, T. Nakagawa, T. Shibayama, K. Uchiya, T. Nikai, K. Ogawa. Comparison of a variable-number tandem-repeat (VNTR) method for typing *Mycobacterium avium* with mycobacterial interspersed repetitive-unit-VNTR and IS*1245* restriction fragment length polymorphism typing. J. Clin. Microbiol. 47 (2009) 2156–2164.

13. K. Arikawa, T. Ichijo, S. Nakajima, Y. Nishiuchi, H. Yano, A. Tamaru, S. Yoshida, F. Maruyama, A. Ota, M. Nasu, D.A. Starkova, I. Mokrousov, O.V. Narvskaya, T. Iwamoto. Genetic relatedness of *Mycobacterium avium* subsp. *hominissuis* isolates from bathrooms of healthy volunteers, rivers, and soils in Japan with human clinical isolates from different geographical areas. Infect. Genet. Evol. 74 (2019) 103923.

14. K. Ichikawa, J. van Ingen, W.J. Koh, D. Wagner, M. Salfinger, T. Inagaki, K.I. Uchiya, T. Nakagawa, K. Ogawa, K. Yamada, T. Yagi. Genetic diversity of clinical *Mycobacterium avium* subsp. *hominissuis* and *Mycobacterium intracellulare* isolates causing pulmonary diseases recovered from different geographical regions. Infect. Genet. Evol. 36 (2015) 250–255.

15. B.R. Imperiale, R.D. Moyano, A.B. DI Giulio, M.A. Romero, M.F. Alvarado Pinedo, M.P. Santangelo, G.E. ravería, N.S. Morcillo, M.I. Romano. Genetic diversity of *Mycobacterium avium* complex strains isolated in Argentina by MIRU-VNTR. Epidemiol. Infect. 145 (2017) 1382–1391.

16. T. Iwamoto, C. Nakajima, Y. Nishiuchi, T. Kato, S. Yoshida, N. Nakanishi, A. Tamaru, Y. Tamura, Y. Suzuki, M. Nasu. Genetic diversity of *Mycobacterium avium* subsp. *hominissuis* strains isolated from humans, pigs, and human living environment. Infect. Genet. Evol. 12 (2012) 846–852.

17. A. Kalvisa, C. Tsirogiannis, I. Silamikelis, G. Skenders, L. Broka, A. Zirnitis, I. Jansone, R. Ranka. MIRU-VNTR genotype diversity and indications of homoplasy in *M. avium* strains isolated from humans and slaughter pigs in Latvia. Infect. Genet. Evol. 43 (2016) 15–21.

18. C. Leão, A. Canto, D. Machado, I.S. Sanches, I. Couto, M. Viveiros, J. Inácio, A. Botelho. Relatedness of *Mycobacterium avium* subspecies *hominissuis* clinical isolates of human and porcine origins assessed by MLVA. Vet. Microbiol. 173 (2014) 92–100.

19. M. Subangkit, T. Yamamoto, M. Ishida, A. Nomura, N. Yasiki, P.E. Sudaryatma, Y. Goto, T. Okabayashi. Genotyping of swine *Mycobacterium avium* subsp. *hominissuis* isolates from Kyushu, Japan. J. Vet. Med. Sci. 81 (2019) 1074–1079.

20. H. Yano, T. Iwamoto, Y. Nishiuchi, C. Nakajima, D.A. Starkova, I. Mokrousov, O. Narvskaya, S. Yoshida, K. Arikawa, N. Nakanishi, K. Osaki, I. Nakagawa, M. Ato, Y. Suzuki, F. Maruyama. Population structure and local adaptation of MAC lung disease agent *Mycobacterium avium* subsp. *hominissuis*. Genome. Biol. Evol. 9 (2017) 2403–2417.

21. H. Yano, H. Suzuki, F. Maruyama, T. Iwamoto. The recombination-cold region as an epidemiological marker of recombinogenic opportunistic pathogen *Mycobacterium avium*. BMC. Genomics. 20 (2019) 752.

22. T. Komatsu, K. Ohya, A. Ota, Y. Nishiuchi, Y. Hirokazu, K. Matsuo, J.O. Odoi, S. Suganuma, K. Sawai, A. Hasebe, T. Asai, T. Yanai, H. Fukushi, T. Wada, S. Yoshida, T. Ito, K. Arikawa, M. Kawai, M. Ato, A.D. Baughn, T. Iwamoto, F. Maruyama. Genomic features of *Mycobacterium avium* subsp. *hominissuis* isolated from pigs in Japan. Gigabyte (2021) gigabyte33.

23. T. Iwamoto, K. Arikawa, C. Nakajima, N. Nakanishi, N. Nishiuchi, Y. Yoshida, A. Tamaru, Y. Tamura, Y. Hoshino, H. Yoo, Y.K. Park, H. Saito, Y. Suzuki. Intra-subspecies sequence variability of the MACPPE12 gene in *Mycobacterium avium* subsp. *hominissuis*. Infect. Genet. Evol. 21 (2014) 479–483.

24. D.A. Starkova, T.F. Otten, I.V. Mokrousov, A.A. Viazovaia, B.I. Vishnevskiĭ, O.V. Narvskaia. Genotypic characteristics of *Mycobacterium avium* subsp. *hominissuis* strains. Genetika. 49 (2013) 1048–1054.

25. R.S. Kaas, P. Leekitcharoenphon, F.M. Aarestrup, O. Lund. Solving the problem of comparing whole bacterial genomes across different sequencing platforms. PLoS. One. 11 (2014) e104984.

26. J. Trifinopoulos, L.T. Nguyen, A. von Haeseler, B.Q. Minh. W-IQ-TREE: a fast online phylogenetic tool for maximum likelihood analysis. Nucleic. Acids. Res. 44 (2016) W232–W235.

27. G. Tonkin-Hill, J.A. Lees, S.D. Bentley, S.D.W. Frost, J. Corander. Fast hierarchical Bayesian analysis of population structure. Nucleic. Acids. Res. 47 (2019) 5539–5549.

28. R. Mostowy, N.J. Croucher, C.P. Andam, J. Corander, W.P. Hanage, P. Marttinen. Efficient inference of recent and ancestral recombination within bacterial populations. Mol. Biol. Evol. 34 (2017) 1167–1182.

29. T. Seemann. Prokka: rapid prokaryotic genome annotation. Bioinformatics. 30 (2014) 2068–2069.

30. A.J. Page, C.A. Cummins, M. Hunt, V.K. Wong, S. Reuter, M.T. Holden, M. Fookes, D. Falush, J.A. Keane, J. Parkhill. Roary: rapid large-scale prokaryote pan genome analysis. Bioinformatics. 31 (2015) 3691–3693.

31. A. Carattoli, E. Zankari, A. García-Fernández, M. Voldby Larsen, O. Lund, L. Villa, F. Møller Aarestrup, H. Hasman. In silico detection and typing of plasmids using PlasmidFinder and plasmid multilocus sequence typing. Antimicrob. Agents. Chemother. 58 (2014) 3895–3903.

32. D. Arndt, J.R. Grant, A. Marcu, T. Sajed, A. Pon, Y. Liang, D.S. Wishart. PHASTER: a better, faster version of the PHAST phage search tool. Nucleic. Acids. Res. 44 (2016) W16–W21.

33. Y. Zhou, Y. Liang, K.H. Lynch, J.J. Dennis, D.S. Wishart. PHAST: a fast phage search tool. Nucleic. Acids. Res. 39 (2011) W347–352.

34. C. Bertelli, M.R. Laird, K.P. Williams, B.Y. Lau, G. Hoad, G.L. Winsor, F.S.L. Brinkman. IslandViewer 4: expanded prediction of genomic islands for larger-scale datasets. Nucleic. Acids. Res. 45 (2017) W30–W35.

35. A. Sanchini, T. Semmler, L. Mao, N. Kumar, F. Dematheis, K. Tandon, V. Peddireddy, N. Ahmed, A. Lewin. A hypervariable genomic island identified in clinical and environmental *Mycobacterium avium* subsp. *hominissuis* isolates from Germany. Int. J. Med. Microbiol. 306 (2016) 495–503.

36. J. Makovcova, M. Slany, V. Babak, I. Slana, P. Kralik. The water environment as a source of potentially pathogenic mycobacteria. J. Water. Health. 12 (2014) 254–263.

37. A. Lahiri, J. Kneisel, I. Kloster, E. Kamal, A. Lewin. Abundance of *Mycobacterium avium* ssp. *hominissuis* in soil and dust in Germany - implications for the infection route. Lett. Appl. Microbiol. 59 (2014) 65–70.

38. G. Ghielmetti, M. Hilbe, U. Friedel, C. Menegatti, L. Bacciarini, R. Stephan, G. Bloemberg. Mycobacterial infections in wild boars (*Sus scrofa*) from Southern Switzerland: diagnostic improvements, epidemiological situation and zoonotic potential. Transbound. Emerg. Dis. 68 (2021) 573–586.

39. J.O. Odoi, K. Ohya, J. Moribe, Y. Takashima, K. Sawai, K. Taguchi, H. Fukushi, T. Wada, S. Yoshida, T. Asai. Isolation and antimicrobial susceptibilities of nontuberculous Mycobacteria from wildlife in Japan. J. Wildl. Dis. 56 (2020) 851–862.

40. S.A. Shoulah, A.M. Oschmann, A. Selim, T. Semmler, C. Schwarz, E. Kamal, F. Hamouda, E. Galila, W. Bitter, A. Lewin. Environmental *Mycobacterium avium* subsp. *hominissuis* have a higher probability to act as a recipient in conjugation than clinical strains. Plasmid. 95 (2018) 28–35.

