## Supplementary for "Unique genomic sequences in a novel *Mycobacterium avium* subsp. *hominissuis* lineage enable fine scale transmission route tracing during pig movement"

[*a]Current Address: Livestock Industry Division, Aichi Agriculture, Forestry and Fisheries, Aichi Prefectural Government, Nagoya, Aichi, Japan

[*b]Current Address: National Institute of Health Sciences, Kawasaki, Kanagawa, Japan

[*c]Current Address: The IDEC Institute, Hiroshima University, Higashi-Hiroshima, Hiroshima, Japan

[#]Corresponding author

**Supplementary Table 1. Information of MAH isolates in this study.**

| Isolate | Farm | Arera | Sampling date | Sampling tissue | Reference |
|---|---|---|---|---|---|
| GM5 | Gifu A | Tokai | 15-Jul-2015 | MEL | 22 |
| GM6 | Gifu A | Tokai | 15-Jul-2015 | MEL | This study |
| GM10 | Gifu A | Tokai | 29-Jul-2015 | MEL | 22 |
| GM11 | Gifu A | Tokai | 29-Jul-2015 | MEL | This study |
| GM12 | Shiga A | Tokai | 05-Aug-2015 | MEL | 22 |
| GM16 | Aichi A | Tokai | 01-Sep-2015 | MEL | 22 |
| GM17 | Aichi B | Tokai | 01-Sep-2015 | MEL | This study |
| GM21 | Aichi B | Tokai | 04-Sep-2015 | MEL | 22 |
| GM23 | Gifu A | Tokai | 07-Sep-2015 | MEL | This study |
| GM24 | Gifu A | Tokai | 07-Sep-2015 | MEL | This study |
| GM25 | Gifu A | Tokai | 07-Sep-2015 | MEL | This study |
| GM32 | Gifu A | Tokai | 05-Oct-2015 | MEL | 22 |
| GM44 | Gifu B | Tokai | 24-Nov-2015 | MEL | 22 |
| OCU467 | Toyama A1 | Hokuriku | 25-Aug-1999 | MEL | 22 |
| OCU468 | Toyama A1 | Hokuriku | 01-Dec-1999 | MEL | 22 |
| OCU469 | Ishikawa B | Hokuriku | 24-Dec-1999 | MEL | 22 |
| OCU470 | Toyama A1 | Hokuriku | 24-Dec-1999 | MEL | 22 |
| OCU471 | Ishikawa B | Hokuriku | 28-Jan-2000 | MEL | 22 |
| OCU472 | Gifu B | Tokai | 04-Feb-2000 | MEL | 22 |
| OCU473 | Ishikawa B | Hokuriku | 08-Feb-2000 | MEL | 22 |
| OCU474 | Ishikawa B | Hokuriku | 22-Dec-2000 | MEL | 22 |
| OCU475 | Toyama D | Hokuriku | 05-Jan-2001 | MEL | 22 |
| OCU476 | Toyama D | Hokuriku | 11-Jan-2001 | MEL | 22 |
| OCU477 | Toyama D | Hokuriku | 11-Jan-2001 | MEL | 22 |
| OCU478 | Toyama A1 | Hokuriku | 18-Jan-2001 | MEL | 22 |
| OCU479 | Ishikawa B | Hokuriku | 23-Jan-2001 | MEL | 22 |
| OCU480 | Ishikawa B | Hokuriku | 26-Jan-2001 | MEL | 22 |
| OCU481 | Toyama D | Hokuriku | 06-Feb-2001 | MEL | 22 |
| OCU482 | Ishikawa B | Hokuriku | 09-Feb-2001 | MEL | 22 |
| OCU483 | Ishikawa B | Hokuriku | 02-Dec-2004 | MEL | 22 |
| OCU484 | Toyama A2 | Hokuriku | 02-Dec-2004 | MEL | 22 |
| OCU485 | Toyama A3 | Hokuriku | 03-Dec-2004 | MEL | 22 |
| OCU486 | Toyama A1 | Hokuriku | 06-Dec-2004 | MEL | 22 |
| Toy194 | Toyama E | Hokuriku | 26-Mar-2018 | Liver | 22 |
| Toy195 | Toyama E | Hokuriku | 26-Mar-2018 | MAL | 22 |
| gifu-1 | Gifu C | Tokai | 11-Mar-2016 | MEL | This study |
| gifu-2 | Gifu C | Tokai | 11-Mar-2016 | MEL | This study |
| gifu-6 | Gifu C | Tokai | 11-Mar-2016 | MEL | This study |
| gifu-7 | Gifu C | Tokai | 11-Mar-2016 | MEL | This study |

| | | | | | |
|---|---|---|---|---|---|
| gifu-8 | Gifu C | Tokai | 11-Mar-2016 | MEL | This study |
| gifu-10 | Gifu C | Tokai | 11-Mar-2016 | MEL | This study |
| gifu-11 | Gifu C | Tokai | 11-Mar-2016 | MEL | This study |
| gifu-34 | Gifu C | Tokai | 22-Mar-2016 | Feces (3 months old ) | This study |
| gifu-41 | Gifu C | Tokai | 22-Mar-2016 | Feces (4 months old ) | This study |
| gifu-50 | Gifu C | Tokai | 22-Mar-2016 | Feces (4 months old ) | This study |
| gifu-51 | Gifu C | Tokai | 22-Mar-2016 | Feces (4 months old ) | This study |
| gifu-53 | Gifu C | Tokai | 22-Mar-2016 | Feces (5 months old) | This study |
| gifu-67 | Gifu C | Tokai | 22-Mar-2016 | Sawdust | This study |
| gifu-77 | Gifu C | Tokai | 22-Mar-2016 | Soil | This study |
| gifu-92 | Gifu C | Tokai | 25-Mar-2016 | MEL | This study |

MEL: mesenteric lymph node.

MAL: mandibular lymph node.

22: Komatsu *et al*., 2021 Gigabyte

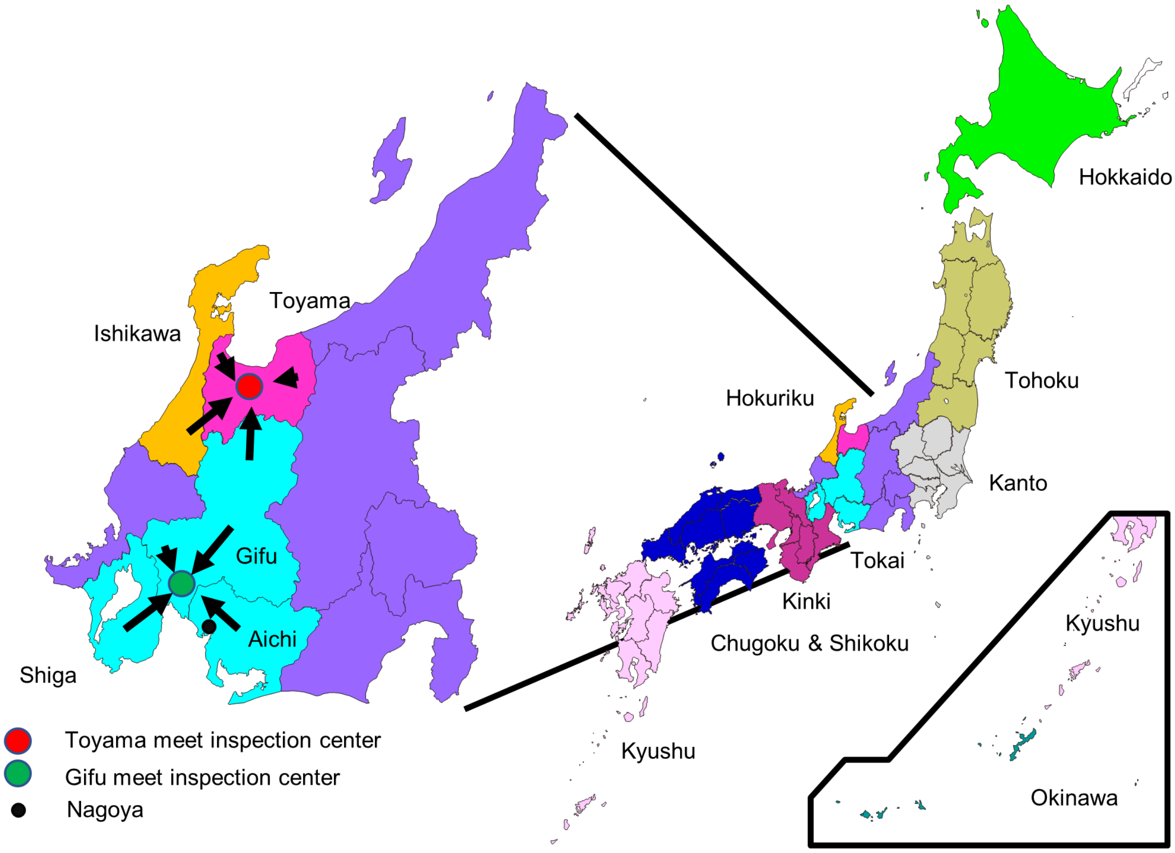


**Supplementary Figure 1. Location of each regions and prefectures in Japan.**

Pigs from Tokai area (southern Gifu, Shiga and Aichi Prefecture) are slaughtered in Gifu meat inspection center. Pigs from Hokuriku area (northern Gifu, Toyama and Ishikawa Prefecture) are slaughtered in Toyama meat inspection center.

### Supplementary Table 2. VNTR profiles of 50 MAH isolates in this study.

| Isolate | MATR and MIRU-VNTR profile | | | | | | | | | | | | | | | | | | | location of farm |
|---|---|---|---|---|---|---|---|---|---|---|---|---|---|---|---|---|---|---|---|---|
| | MATR -1 | MATR -2 | MATR -3 | MATR -4 | MATR -5 | MATR -6 | MATR -7 | MATR -8 | MATR -9 | MATR -11 | MATR -12 | MATR -13 | MATR -14 | MATR -15 | MATR -16 | | | | | |
| | | MIRU-292 | MIRU-X3 | | | | | | MIRU-10 | | | | | | | MIRU -7 | MIRU -25 | MIRU -32 | MIRU -47 | |
| OCU467 | 2 | 2 | 3 | 2 | 3 | 3 | 2 | 2 | 2 | 0 | 3 | 2 | 3 | 2 | 0 | 1 | 2 | 9 | 2 | Toyama |
| OCU468 | 1 | 2 | 4 | 2 | 3 | 3 | 3 | 3 | 2 | 5 | 3 | 2 | 4 | 4 | 3 | 1 | 2 | 9 | 2 | Toyama |
| OCU469 | 2 | 0 | 3 | 3 | 3 | 3 | 4 | 5 | 2 | 3 | 3 | 2 | 3 | 2 | 3 | 1 | 3 | 9 | 2 | Ishikawa |
| OCU470 | 1 | 2 | 4 | 2 | 3 | 3 | 3 | 3 | 2 | 5 | 3 | 2 | 4 | 4 | 3 | 1 | 3 | 9 | 2 | Toyama |
| OCU471 | 2 | 2 | 2 | 1 | 3 | 3 | 4 | 1 | 1 | 4 | 3 | 2 | 3 | 3 | 3 | 1 | 2 | 9 | 2 | Ishikawa |
| OCU472 | 1 | 2 | 4 | 2 | 3 | 3 | 3 | 3 | 2 | 5 | 3 | 2 | 4 | 4 | 3 | 1 | 2 | 9 | 2 | Gifu |
| OCU473 | 2 | 2 | 3 | 2 | 3 | 3 | 3 | 4 | 2 | 3 | 3 | 2 | 5 | 4 | 3 | 1 | 3 | 9 | 2 | Ishikawa |
| OCU474 | 2 | 2 | 2 | 1 | 3 | 3 | 4 | 1 | 1 | 0 | 3 | 2 | 3 | 4 | 2 | 1 | 2 | 9 | 2 | Ishikawa |
| OCU475 | 1 | 2 | 4 | 2 | 3 | 3 | 3 | 3 | 2 | 5 | 3 | 2 | 4 | 4 | 3 | 1 | 2 | 9 | 2 | Toyama |
| OCU476 | 1 | 2 | 4 | 2 | 3 | 3 | 3 | 3 | 2 | 5 | 3 | 2 | 4 | 4 | 3 | 1 | 2 | 9 | 2 | Toyama |
| OCU477 | 2 | 3 | 2 | 1 | 1 | 3 | 4 | 1 | 1 | 4 | 3 | 2 | 2 | 4 | 2 | 1 | 2 | 7 | 2 | Toyama |
| OCU478 | 2 | 2 | 3 | 2 | 3 | 3 | 2 | 2 | 2 | 3 | 3 | 2 | 3 | 2 | 2 | 1 | 2 | 7 | 2 | Toyama |
| OCU479 | 1 | 2 | 4 | 2 | 3 | 3 | 3 | 2 | 2 | 5 | 3 | 2 | 4 | 4 | 2 | 1 | 2 | 7 | 2 | Ishikawa |
| OCU480 | 2 | 2 | 2 | 1 | 3 | 3 | 4 | 1 | 1 | 4 | 3 | 2 | 3 | 4 | 2 | 1 | 3 | 9 | 2 | Ishikawa |
| OCU481 | 1 | 2 | 3 | 2 | 3 | 3 | 3 | 2 | 1 | 5 | 3 | 2 | 3 | 2 | 2 | 1 | 2 | 9 | 2 | Toyama |
| OCU482 | 1 | 2 | 3 | 2 | 3 | 3 | 3 | 2 | 1 | 5 | 3 | 2 | 0 | 2 | 2 | 1 | 2 | 9 | 2 | Ishikawa |
| OCU483 | 2 | 1 | 4 | 1 | 2 | 1 | 4 | 1 | 2 | 4 | 3 | 2 | 2 | 2 | 0 | 1 | 2 | 9 | 2 | Ishikawa |
| OCU484 | 2 | 3 | 4 | 2 | 3 | 3 | 4 | 4 | 2 | 2 | 3 | 2 | 2 | 2 | 3 | 1 | 3 | 9 | 2 | Toyama |
| OCU485 | 2 | 2 | 3 | 2 | 3 | 3 | 2 | 2 | 2 | 3 | 3 | 2 | 2 | 2 | 3 | 1 | 3 | 9 | 2 | Toyama |
| OCU486 | 2 | 1 | 2 | 1 | 1 | 3 | 4 | 1 | 1 | 4 | 3 | 2 | 2 | 4 | 3 | 1 | 2 | 9 | 2 | Toyama |
| Toy-194 | 0 | 3 | 5 | 2 | 3 | 3 | 3 | 2 | 2 | 4 | 3 | 2 | 4 | 4 | 3 | 1 | 2 | 9 | 2 | Toyama |
| Toy-195 | 2 | 3 | 5 | 2 | 3 | 3 | 4 | 2 | 2 | 2 | 3 | 2 | 2 | 4 | 3 | 1 | 2 | 9 | 2 | Toyama |
| gifu-1 | 2 | 2 | 2 | 2 | 2 | 3 | 4 | 1 | 2 | 2 | 3 | 2 | 2 | 4 | 1 | 1 | 3 | 8 | 3 | Gifu |
| gifu-2 | 2 | 2 | 5 | 2 | 3 | 1 | 4 | 1 | 2 | 2 | 3 | 2 | 2 | 2 | 2 | 1 | 2 | 8 | 2 | Gifu |
| gifu-6 | 2 | 2 | 2 | 2 | 2 | 3 | 4 | 1 | 2 | 4 | 3 | 2 | 2 | 4 | 1 | 1 | 3 | 8 | 3 | Gifu |
| gifu-7 | 2 | 2 | 2 | 2 | 3 | 1 | 4 | 1 | 1 | 4 | 3 | 2 | 2 | 4 | 1 | 1 | 4 | 8 | 3 | Gifu |
| gifu-8 | 2 | 2 | 6 | 2 | 2 | 1 | 4 | 1 | 2 | 4 | 3 | 2 | 2 | 4 | 2 | 1 | 3 | 8 | 3 | Gifu |

| | | | | | | | | | | | | | | | | | | | | |
|---|---|---|---|---|---|---|---|---|---|---|---|---|---|---|---|---|---|---|---|---|
| gifu-10 | 2 | 2 | 2 | 2 | 2 | 3 | 4 | 1 | 2 | 4 | 3 | 2 | 2 | 4 | 1 | 1 | 3 | 8 | 3 | Gifu |
| gifu-11 | 2 | 2 | 2 | 2 | 2 | 3 | 4 | 1 | 2 | 4 | 3 | 2 | 2 | 4 | 1 | 1 | 3 | 8 | 3 | Gifu |
| gifu-34 | 2 | 2 | 2 | 2 | 3 | 1 | 4 | 1 | 1 | 4 | 3 | 2 | 2 | 4 | 1 | 1 | 2 | 8 | 3 | Gifu |
| gifu-41 | 2 | 2 | 2 | 2 | 2 | 3 | 4 | 1 | 2 | 4 | 3 | 2 | 2 | 4 | 1 | 1 | 3 | 8 | 3 | Gifu |
| gifu-50 | 1 | 2 | 4 | 2 | 3 | 3 | 3 | 3 | 2 | 5 | 3 | 2 | 4 | 4 | 2 | 1 | 2 | 8 | 2 | Gifu |
| gifu-51 | 2 | 2 | 5 | 2 | 3 | 1 | 4 | 2 | 2 | 6 | 3 | 2 | 2 | 2 | 3 | 1 | 3 | 8 | 3 | Gifu |
| gifu-53 | 2 | 2 | 2 | 2 | 3 | 1 | 4 | 1 | 1 | 4 | 3 | 2 | 2 | 4 | 1 | 1 | 4 | 8 | 3 | Gifu |
| gifu-67 | 2 | 2 | 2 | 2 | 2 | 3 | 4 | 1 | 2 | 4 | 3 | 2 | 2 | 4 | 1 | 1 | 3 | 8 | 3 | Gifu |
| gifu-77 | 2 | 2 | 2 | 1 | 2 | 1 | 4 | 1 | 1 | 4 | 3 | 2 | 2 | 2 | 1 | 1 | 4 | 8 | 3 | Gifu |
| gifu-92 | 2 | 2 | 2 | 1 | 2 | 1 | 4 | 1 | 1 | 4 | 3 | 2 | 2 | 2 | 1 | 1 | 4 | 8 | 3 | Gifu |
| GM5 | 2 | 0 | 3 | 3 | 2 | 3 | 4 | 4 | 2 | 2 | 3 | 2 | 2 | 2 | 3 | 1 | 4 | 8 | 3 | Gifu |
| GM6 | 2 | 0 | 3 | 3 | 2 | 3 | 4 | 4 | 2 | 2 | 3 | 2 | 2 | 2 | 3 | 1 | 4 | 8 | 3 | Gifu |
| GM10 | 2 | 4 | 3 | 1 | 2 | 2 | 1 | 3 | 4 | 3 | 3 | 2 | 2 | 1 | 2 | 1 | 1 | 8 | 3 | Gifu |
| GM11 | 2 | 4 | 3 | 1 | 2 | 2 | 1 | 3 | 4 | 3 | 3 | 2 | 2 | 1 | 2 | 1 | 1 | 8 | 3 | Gifu |
| GM12 | 2 | 4 | 3 | 1 | 2 | 2 | 1 | 3 | 4 | 3 | 3 | 2 | 2 | 1 | 2 | 1 | 1 | 8 | 3 | Shiga |
| GM16 | 5 | 3 | 3 | 2 | 1 | 0 | 1 | 1 | 2 | 2 | 3 | 2 | 2 | 2 | 3 | 1 | 2 | 8 | 3 | Aichi |
| GM17 | 1 | 3 | 3 | 2 | 3 | 1 | 3 | 2 | 5 | 2 | 2 | 2 | 1 | 2 | 2 | 1 | 2 | 8 | 3 | Aichi |
| GM21 | 2 | 4 | 3 | 1 | 2 | 2 | 1 | 3 | 4 | 3 | 3 | 2 | 2 | 1 | 2 | 1 | 1 | 8 | 3 | Aichi |
| GM23 | 2 | 4 | 3 | 1 | 2 | 2 | 1 | 3 | 4 | 3 | 3 | 2 | 2 | 1 | 2 | 1 | 1 | 8 | 3 | Aichi |
| GM24 | 2 | 4 | 3 | 1 | 2 | 2 | 1 | 3 | 4 | 3 | 3 | 2 | 2 | 1 | 2 | 1 | 1 | 8 | 3 | Gifu |
| GM25 | 2 | 4 | 3 | 1 | 2 | 2 | 1 | 3 | 4 | 3 | 3 | 2 | 2 | 1 | 2 | 1 | 1 | 8 | 3 | Gifu |
| GM32 | 2 | 4 | 3 | 1 | 2 | 2 | 1 | 3 | 4 | 3 | 3 | 2 | 2 | 1 | 2 | 1 | 1 | 8 | 3 | Gifu |
| GM44 | 2 | 3 | 4 | 2 | 3 | 3 | 3 | 2 | 5 | 4 | 3 | 2 | 1 | 2 | 2 | 1 | 2 | 8 | 2 | Gifu |

**Supplementary Table 3. Information of MAH strains or isolates used for VNTR analysis in this study.**

| Nation | Region in Japan | No. of strains | | | | | | Reference** |
|---|---|---|---|---|---|---|---|---|
| | | human | | | Bathroom | Natural environment | Pig | |
| | | HIV | | | | | | |
| | | negative | positive | unknown | | | | |
| Japan | Hokkaido | | | 31 | 3 | | 5 | 13, 16 |
| | Tohoku | | | | 3 | | | 13 |
| | Kanto | | | | 15 | | | 13 |
| | Chubu | 97 | | 94 | 4 | 7* | 43* | 4, 13, 14 |
| | Kinki | 93 | | | 60 | 22 | 70 | 13, 16 |
| | Chugoku & Shikoku | | | | 8 | | | 13 |
| | Kyushu | | | | | | 36 | 19 |
| | Okinawa | | | | | | 19 | 4 |
| | Unknown | | 28 | | | | | 4 |
| Korea | | 77 | | 98 | | | | 14, 23 |
| USA | | | | 32 | | | | 14 |
| Europe | | | | 37 | | | | 14, 24 |
| Russia | | 65 | 25 | | | | | 23 |

total: 972 strains or isolates

*: Isolates in this study

**: The numbers of references are the same as the manuscript and are as follows.

4: Adachi *et al*., 2016 J Med Microbiol

13: Arikawa *et al*., 2019 Infect Genet Evol

14: Ichikawa *et al*., 2015 Infect Genet Evol

16: Iwamoto *et al*., 2012 Infect Genet Evol

19: Subangkit *et al*., 2019 J vet med sci

23: Iwamoto *et al*., 2014 Infect Genet Evol

24: Starkova *et al*., 2013 Genetika

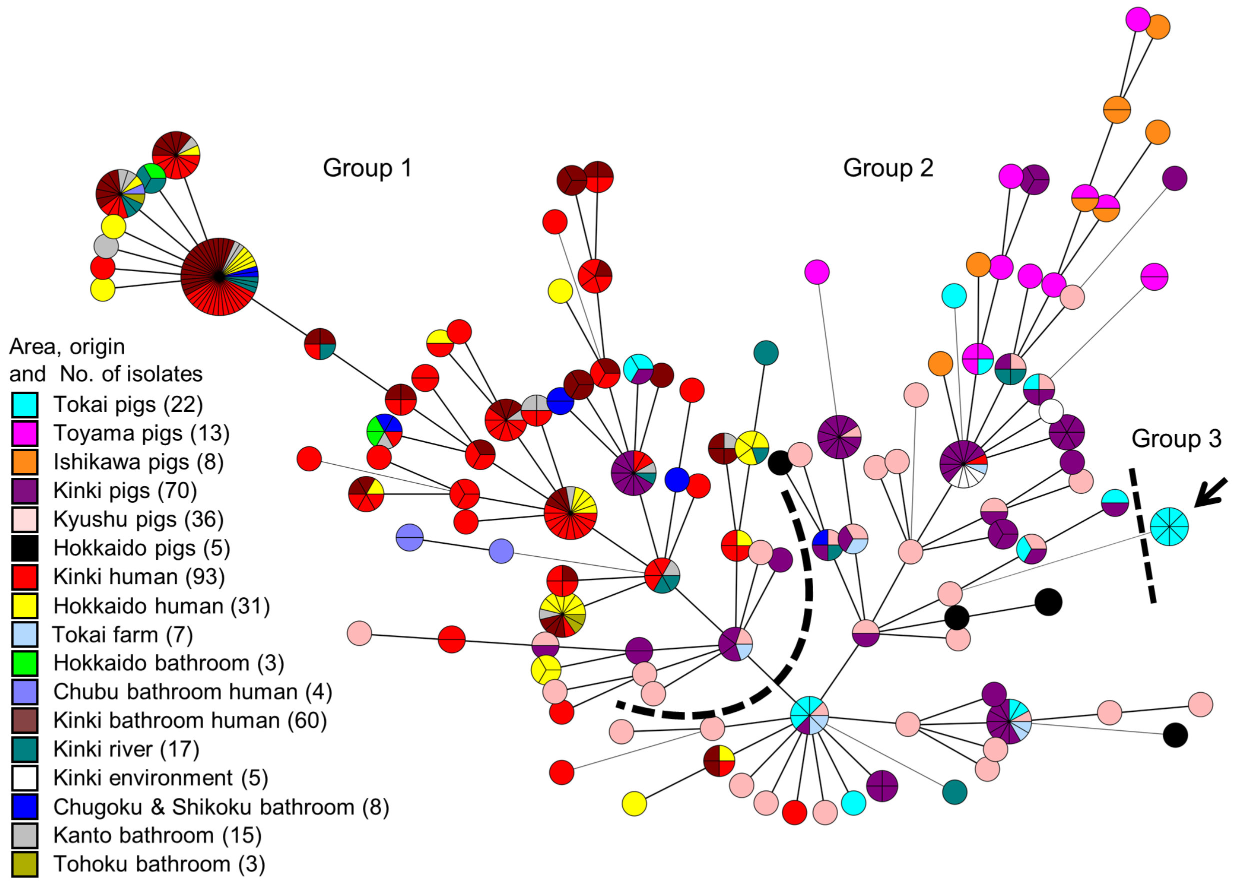


**Supplementary Figure 2. Minimum spanning tree based on 7 loci Mycobacterial Interspersed Repetitive Unit VNTR genotyping of MAH isolates in Japan.** Circles indicate different VNTR profiles. The size of each circle depended on the number of isolates sharing the same profiles. New lineages indicated by an arrow were clustered and consisted of isolates GM10, GM11, GM12, GM21, GM23, GM24, GM25, GM32.

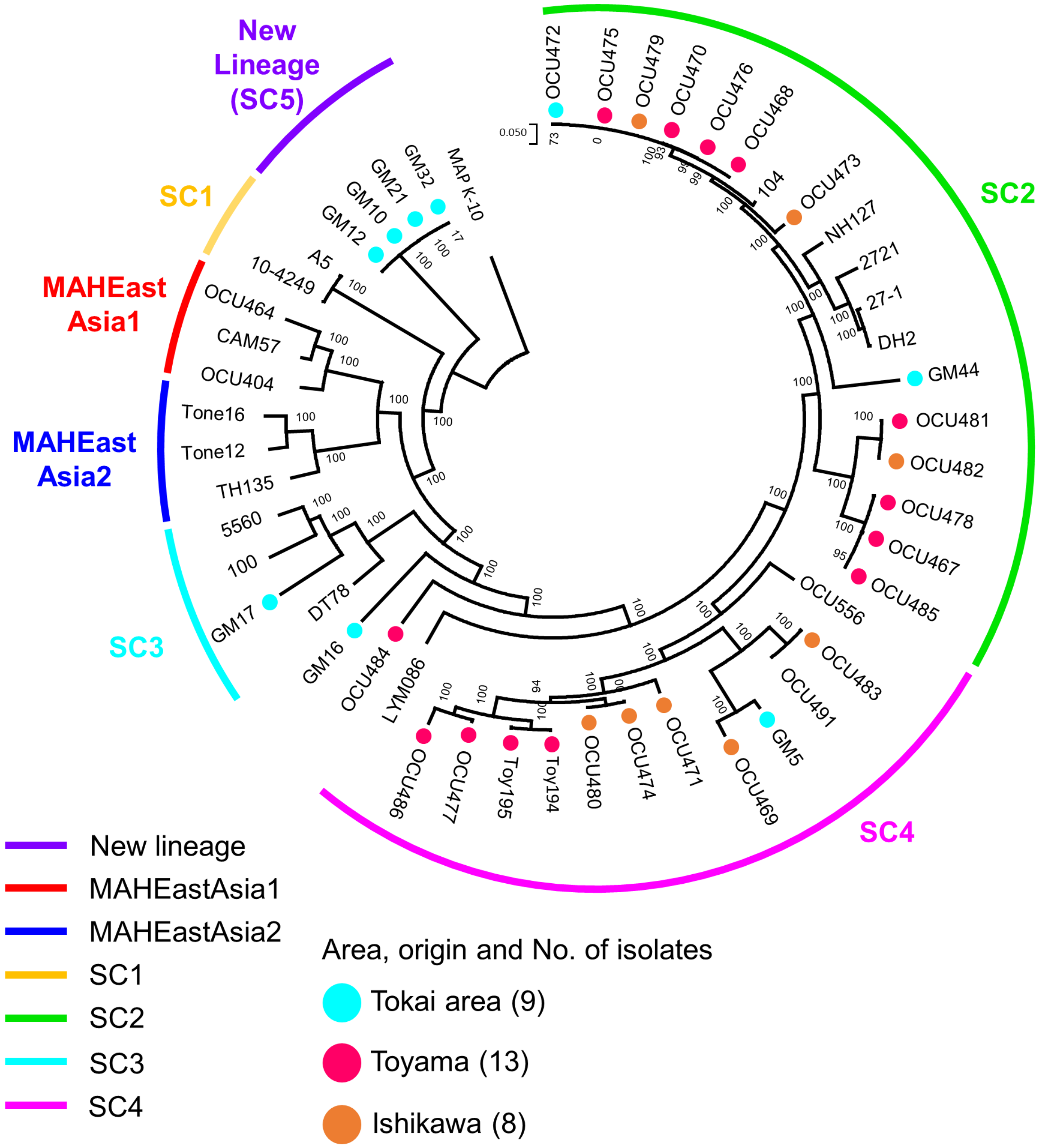


**Supplementary Figure 3. Phylogenetic tree based on draft genome sequences generated by CSIphylogeny.** Maximum likelihood tree was generated by CSI phylogeny version 1.4 (25) and was visualized via MEGA 7.0. The area and Prefecture of isolates in this study were indicated by the color dots. Lineage classification except for new lineage was based on Yano *et al* (21).

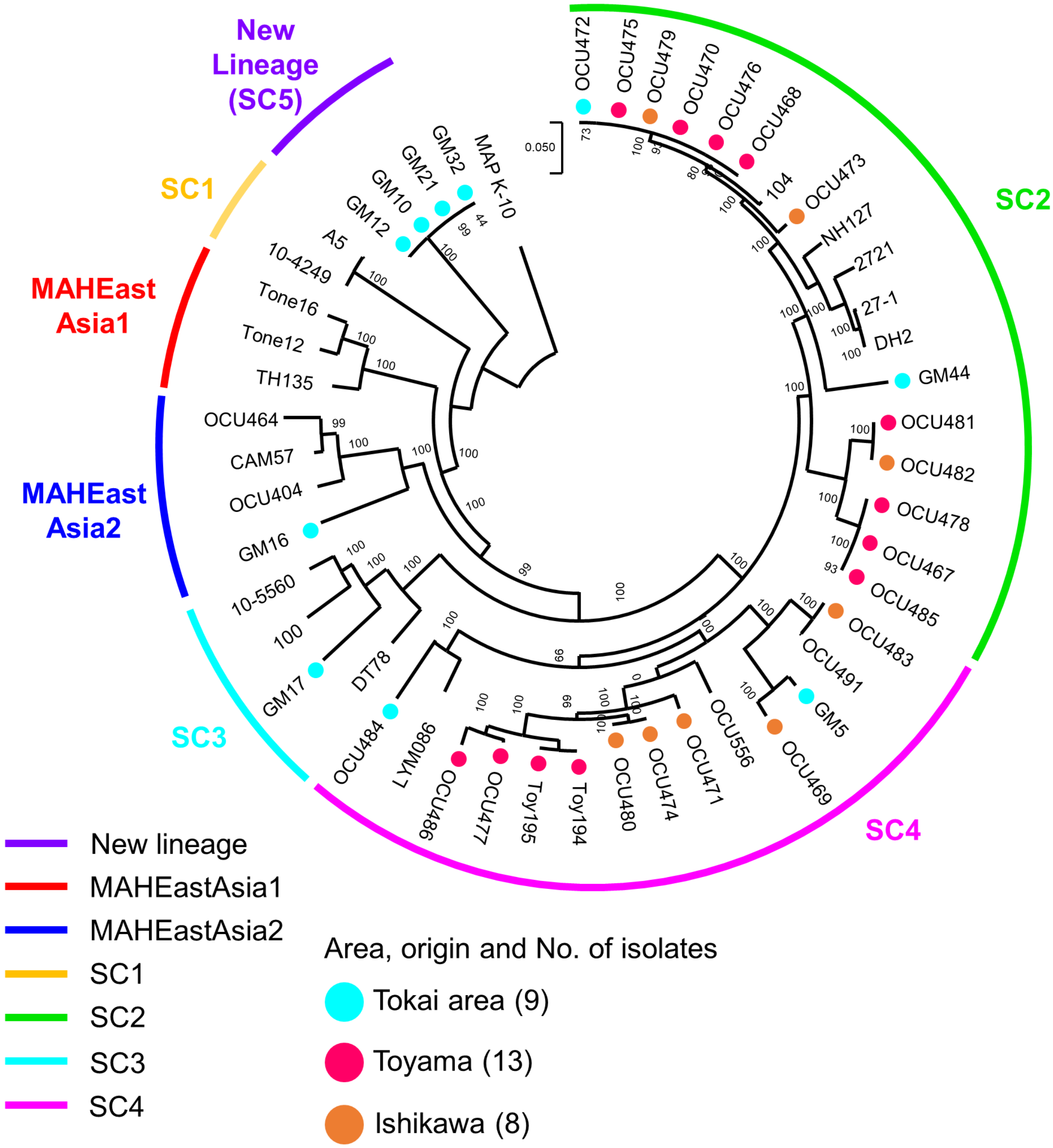


**Supplementary Figure 4. Phylogenetic tree based on draft genome sequences generated by IQ-TREE.** Maximum likelihood tree was generated by IQ-TREE (26) and visualized via MEGA 7.0. The Area and Prefecture of isolates in this study were indicated by the color dots. Lineage classification except for new lineage was based on Yano *et al* (21).

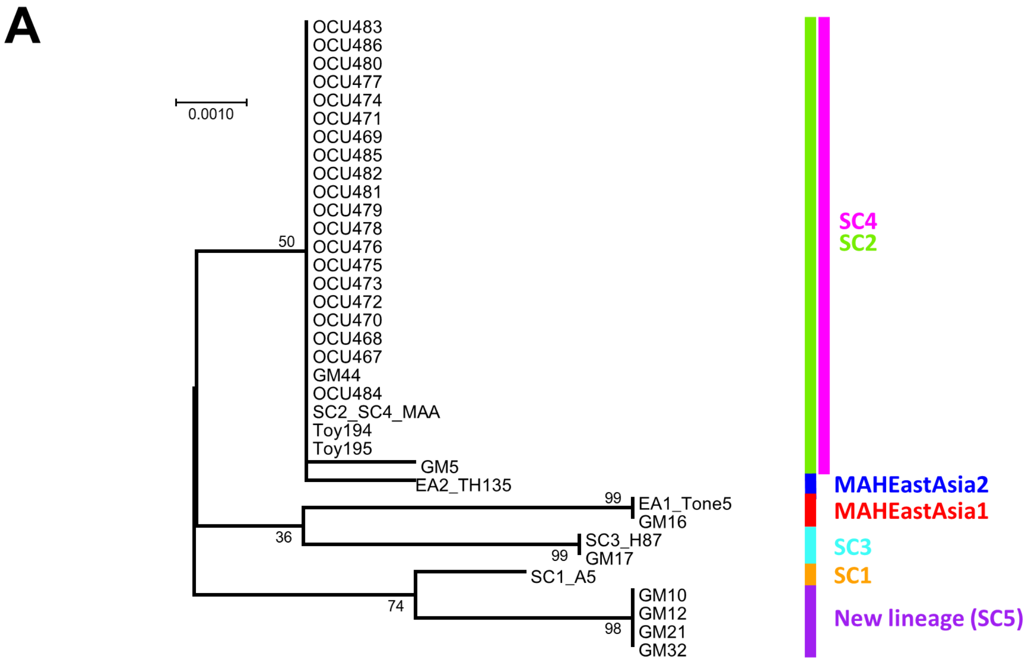


**B**

| | | SNP position | 36 | 72 | 157 | 198 | 231 | 240 | 267 | 288 | 351 | 552 | 564 | 570 | 705 | 708 | 816 | 888 | 897 | 900 | 918 | 1077 | 1140 | 1239 | 1269 | 1317 |
|---|---|---|---|---|---|---|---|---|---|---|---|---|---|---|---|---|---|---|---|---|---|---|---|---|---|---|
| Reference | | EA1 (Tone5) | C | G | T | T | C | T | C | G | T | C | C | A | C | A | T | C | T | C | C | G | G | G | A | A |
| Reference | | EA2 (TH135) | C | G | T | T | C | C | C | A | C | C | G | G | C | A | T | C | C | G | G | G | G | G | G | G |
| Reference | | SC1 (A5) | C | T | T | C | T | C | G | G | C | C | G | G | C | A | T | C | C | G | C | C | C | A | A | G |
| Reference | | SC2/SC4/MAA | C | G | T | T | C | C | C | A | C | C | G | G | C | A | T | C | C | G | G | G | G | A | A | G |
| Reference | | SC3 (H87) | C | G | T | T | C | T | C | G | C | T | G | G | G | G | T | T | C | G | C | G | G | G | G | G |
| Reference | | MAS (ATCC49884) | C | G | T | T | C | C | C | A | C | C | G | G | C | A | C | C | C | G | G | G | G | A | A | G |
| Reference | | MAP (K10) | - | G | C | C | C | C | G | G | T | C | G | G | C | A | T | C | C | G | C | C | C | A | A | G |
| This study | New lineage (SC5) | GM10 | T | G | T | C | C | C | G | G | T | C | G | G | C | A | T | C | T | C | C | C | C | A | A | G |
| This study | New lineage (SC5) | GM12 | T | G | T | C | C | C | G | G | T | C | G | G | C | A | T | C | T | C | C | C | C | A | A | G |
| This study | New lineage (SC5) | GM21 | T | G | T | C | C | C | G | G | T | C | G | G | C | A | T | C | T | C | C | C | C | A | A | G |
| This study | New lineage (SC5) | GM32 | T | G | T | C | C | C | G | G | T | C | G | G | C | A | T | C | T | C | C | C | C | A | A | G |
| This study | SC2 | GM44, OCU467 OCU468, OCU470 OCU472, OCU473 OCU475, OCU476 OCU478, OCU479 OCU481, OCU482 OCU485 | C | G | T | T | C | C | C | A | C | C | G | G | C | A | T | C | C | G | G | G | G | A | A | G |
| This study | SC3 | GM17 | C | G | T | T | C | T | C | G | C | T | G | G | G | G | T | T | C | G | C | G | G | G | G | G |
| This study | SC4 | GM5 | C | G | T | T | C | C | C | A | C | C | C | A | C | A | T | C | C | G | G | G | G | A | A | G |
| This study | SC4 | GM16 | C | G | T | T | C | T | C | G | T | C | C | A | C | A | T | C | T | C | C | G | G | G | A | A |
| This study | SC4 | OCU469, OCU471 OCU474, OCU477 OCU480, OCU483 OCU484, OCU486 Toy194, Toy195 | C | G | T | T | C | C | C | A | C | C | G | G | C | A | T | C | C | G | G | G | G | A | A | G |

**Supplementary Figure 5. Allelic variants in *cinA* gene of pig MAH.** A) Maximum likelihood tree generated by MEGA 7.0. Bootstrap values were calculated by 1,000 replications. The scale bar indicates genetic distance of each strain. Lineage classification except for new lineage was based on Yano *et al* (21). B) SNP cites in pig

MAH. New lineage shared the same SNP patterns, which were highlighted by gray. Some variations were detected in SC4. Reference SNP patterns were referred from Yano *et al* (21).

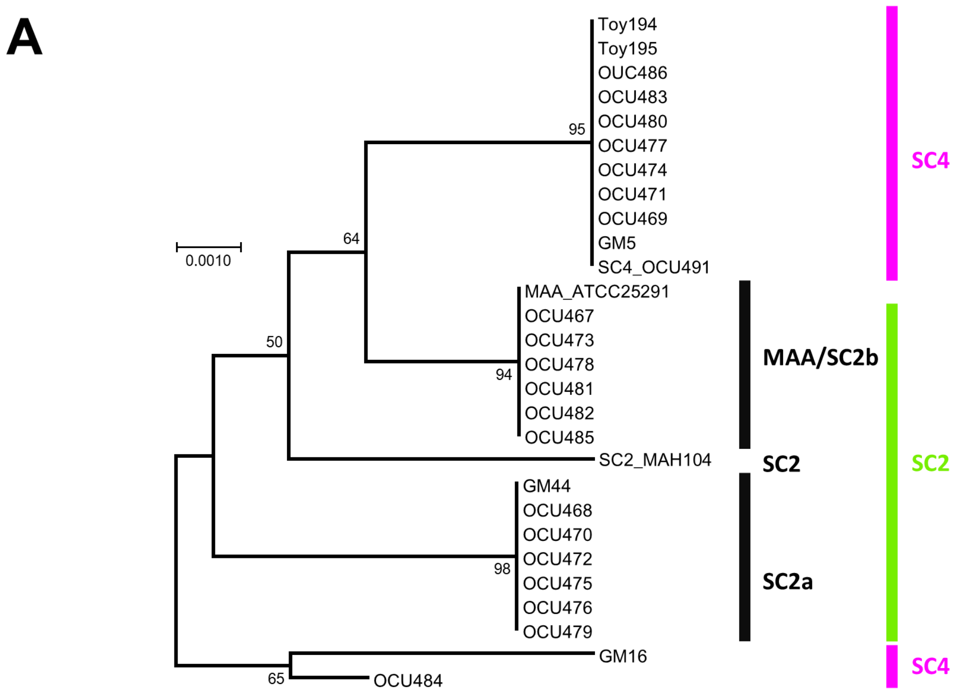


**B**

| | | SNP position | 25 | 71 | 165 | 168 | 298 | 336 | 387 | 393 | 432 | 441 | 495 | 592 | 654 | 664 | 669 | 705 | 789 | 852 | 855 | 868 |
|---|---|---|---|---|---|---|---|---|---|---|---|---|---|---|---|---|---|---|---|---|---|---|
| Reference | | SC2 (Mah104) | C | T | T | C | A | G | G | G | C | C | G | C | C | T | A | C | A | A | C | G |
| Reference | | SC4 (OCU491) | C | C | C | T | G | G | G | A | C | C | C | C | C | T | A | C | G | C | C | G |
| Reference | | MAA (ATCC25291) | C | C | T | C | G | G | G | G | A | C | C | C | A | T | A | C | G | C | C | G |
| This study | New lineage (SC5) | GM10 | C | C | C | T | G | C | G | A | C | C | C | C | A | T | A | C | G | C | C | G |
| This study | New lineage (SC5) | GM12 | C | C | C | T | G | C | G | A | C | C | C | C | A | T | A | C | G | C | C | G |
| This study | New lineage (SC5) | GM21 | C | C | C | T | G | C | G | A | C | C | C | C | A | T | A | C | G | C | C | G |
| This study | New lineage (SC5) | GM32 | C | C | C | T | G | C | G | A | C | C | C | C | A | T | A | C | G | C | C | G |
| This study | SC2 / SC2a | GM44, OCU468 OCU470, OCU472 OCU475, OCU476 OCU479 | T | C | C | C | G | G | G | G | C | C | G | T | C | C | G | C | G | C | C | G |
| This study | SC2 / SC2b | OCU467, OCU473 OCU478, OCU481 OCU482, OCU485 | C | C | T | C | G | G | G | G | A | C | C | C | A | T | A | C | G | C | C | G |
| This study | SC3 | GM17 | C | T | T | C | G | G | G | G | C | C | C | C | C | T | A | T | G | C | C | G |
| This study | SC4 | GM5, OCU469 OCU471, OCU474 OCU477, OCU480 OCU483, OCU486 Toy194, Toy195 | C | C | C | T | G | G | G | A | C | C | C | C | C | T | A | C | G | C | C | G |
| This study | SC4 | GM16 | C | C | T | C | G | G | A | G | C | G | G | T | C | T | G | T | G | A | G | G |
| This study | SC4 | OCU484 | C | C | T | C | G | G | A | G | C | G | G | T | C | T | A | C | G | C | C | C |

**Supplementary Figure 6. Allelic variants in *sugA* gene of pig MAH.** A) Maximum likelihood tree generated by MEGA 7.0. Bootstrap values were calculated by 1,000 replications. The scale bar indicates genetic distance of each strain. Lineage classification except for new lineage was based on Yano *et al* (21). B) SNP cites in pig

MAH. New lineage shared the same SNP patterns, which were highlighted by gray. Two type of SNP patterns were detected in SC2. Reference SNP patterns were referred from Yano *et al* (21).

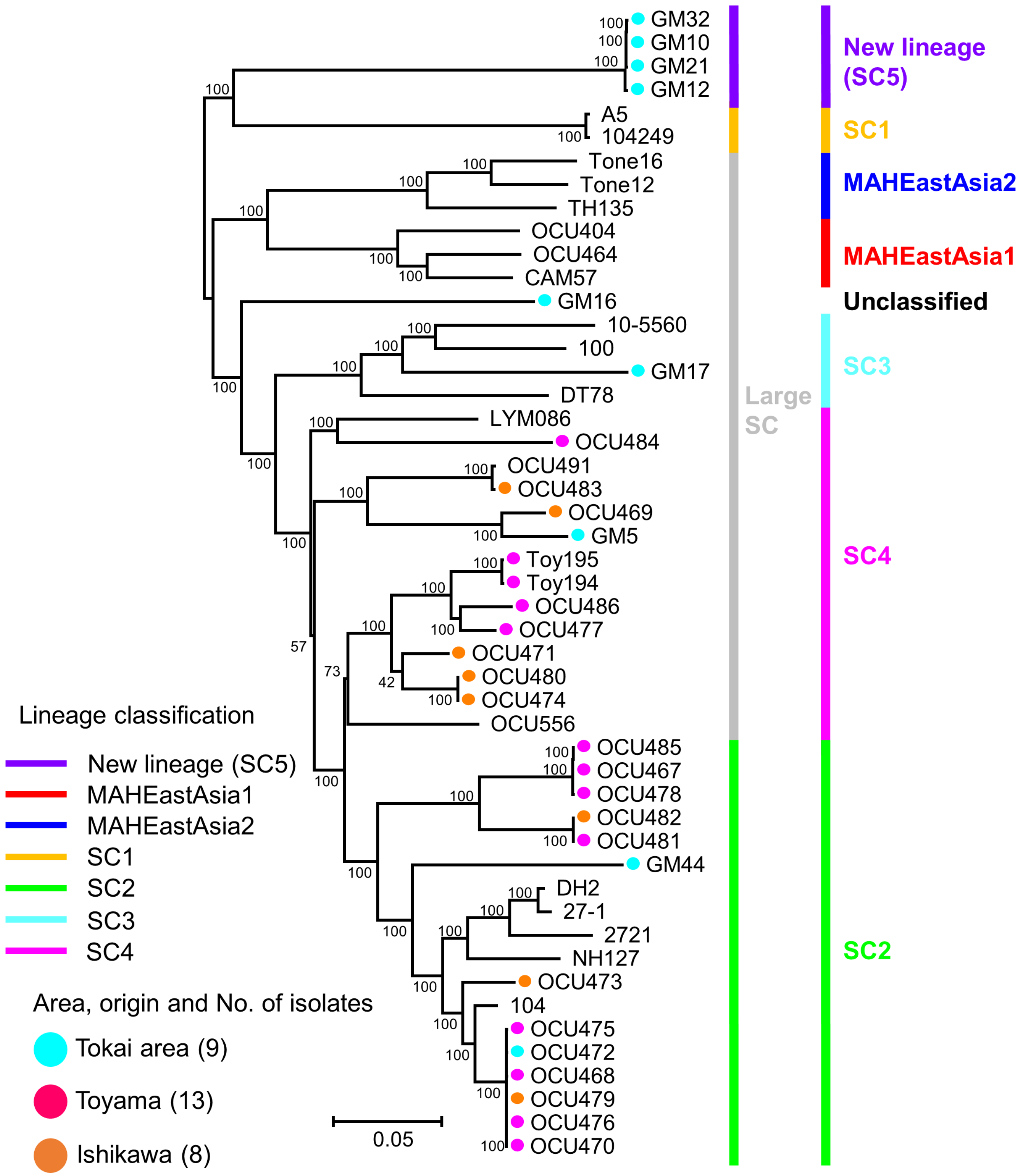


**Supplementary Figure 7. Lineage classification by fastBAPS.** Lineage classification was predicted by fastBAPS (27). Bootstrap values were calculated by 1,000 replications. The scale bar indicates genetic distance of each strain. Lineage classification in the right side except for new lineage was based on Yano *et al* (21). Neighbor joining tree was generated by via MEGA 7.0 using alignment file obtained from CSI phylogeny version 1.4 (25).

### Supplementary Table 4. Uniquely present or absent genes in new lineage (SC5).

| Absent/ Present* | Annotated protein | Genes on the genomic island (GI) / phage (Ph) | | | | BLAST search result | | | | |
|---|---|---|---|---|---|---|---|---|---|---|
| | | GM10 | GM12 | GM21 | GM32 | Strain | Coverage | Identity | E-value | Accession |
| Absent gene (6) | hypothetical protein | - | - | - | - | | | | | |
| | Serine/threonine-protein kinase PknL | - | - | - | - | | | | | |
| | hypothetical protein | - | - | - | - | | | | | |
| | hypothetical protein | - | - | - | - | | | | | |
| | hypothetical protein | - | - | - | - | | | | | |
| | hypothetical protein | - | - | - | - | | | | | |
| Present gene (97) | hypothetical protein | - | - | - | - | MAH strain OCU464[a] | 99% | 100% | 0.00E+00 | CP009360 |
| | hypothetical protein | - | - | - | - | MAH strain 101115[a] | 99% | 100% | 2.00E-94 | CP040255 |
| | hypothetical protein | - | - | - | - | *Chloropicon primus* strain CCMP1205 | 2% | 88.37% | 0.031 | CP031034 |
| | IS*481* family transposase IS*Mav5* | GI | GI | GI | GI | MAH strain mc2 2500[a] | 100% | 99.02% | 0.00E+00 | CP036220 |
| | hypothetical protein | - | - | GI | GI | *M. canettii* CIPT 140010059 | 100% | 76.70% | 7.00E-127 | HE572590 |
| | hypothetical protein | - | - | - | - | *M. colombiense* CECT 3035 | 83% | 87.17% | 3.00E-56 | CP020821 |
| | hypothetical protein | GI | GI | GI | GI | *M. sp.* KMS | 100% | 84.36% | 0.00E+00 | CP000518 |
| | hypothetical protein | GI | - | - | GI | *M. sp.* KMS | 98% | 74.24% | 5.00E-42 | CP000518 |
| | hypothetical protein | GI | GI | GI | GI | *M. sp.* KMS | 75% | 82.45% | 0.00E+00 | CP000518 |
| | hypothetical protein | GI | GI | GI | GI | *M. branderi* JCM 12687 | 97% | 83.72% | 0.00E+00 | AP022606 |
| | hypothetical protein | GI | GI | GI | GI | Mycobacterium phage Dori | 75% | 67.58% | 9.00E-54 | NC_023703 |
| | Tyrosine recombinase XerC | GI | GI | GI | GI | *M. chimaera* strain AUSMDU00007395 | 98% | 68.74% | 4.00E-146 | CP045963 |
| | Serine recombinase PinR | GI | GI | GI | GI | *Rhodococcus biphenylivorans* strain TG9 | 67% | 69.21% | 6.00E-39 | CP022208 |
| | hypothetical protein | GI | GI | GI | GI | *M. chimaera* strain AUSMDU00007395 | 96% | 83.67% | 5.00E-66 | CP045963 |
| | hypothetical protein | GI | GI | GI | GI | *Mycolicibacterium alvei* JCM 12272 | 25% | 71.26% | 2.00E-71 | AP022565 |

| | | | | | | | | | |
|---|---|---|---|---|---|---|---|---|---|
| hypothetical protein | GI | GI | GI | GI | PREDICTED: *Monopterus albus* | 13% | 93.75% | 0.65 | XM_020592160 |
| hypothetical protein | GI | GI | GI | GI | *Hordeum vulgare* subsp. *vulgare* | 7% | 81.36% | 0.017 | AK376348 |
| hypothetical protein | GI | GI | GI | GI | *M. marinum* M | 79% | 71.65% | 1.00E-17 | CP000854 |
| hypothetical protein | GI | GI | GI | GI | *M. intracellulare* ATCC 13950 | 94% | 73.97% | 0.00E+00 | CP003322 |
| hypothetical protein | GI | GI | GI | GI | *Streptomyces sp.* RTd22 | 9% | 100% | 0.002 | CP015726 |
| hypothetical protein | GI | GI | GI | GI | *M. marinum* strain 1218R | 75% | 84.69% | 3.00E-20 | CP025779 |
| hypothetical protein | GI | - | GI | GI | MAH strain MAC109[a] | 100% | 98.00% | 1.00E-64 | CP029332 |
| IS*481* family transposase IS*3514* | GI | GI | GI | GI | Mycolicibacterium helvum JCM 30396 | 100% | 79.17% | 0.00E+00 | AP022596 |
| hypothetical protein | GI | GI | GI | GI | *M. shottsii* JCM 12657 | 17% | 76.40% | 2.00E-06 | AP022572 |
| Tyrosine recombinase XerC | GI | GI | GI | GI | *M. saskatchewanense* JCM 13016 | 99% | 74.67% | 1.00E-117 | AP022573 |
| hypothetical protein | GI | GI | GI | GI | *M. conspicuum* JCM 14738 | 50% | 74.44% | 4.00E-11 | AP022613 |
| hypothetical protein | GI | GI | GI | GI | *Rathayibacter sp.* VKM Ac-2804 | 23% | 73.73% | 2.00E-08 | CP047420 |
| hypothetical protein | GI | GI | GI | GI | *Rhodobacteraceae bacterium* SH-1 | 30% | 84.62% | 1.1 | CP027665 |
| Putative prophage phiRv2 integrase | GI | GI | GI | GI | *M. sp.* JS623 | 96% | 74.40% | 0.00E+00 | CP003078 |
| hypothetical protein | - | - | - | - | MAA strain DSM 44156 | 95% | 80.62% | 0.00E+00 | CP046507 |
| hypothetical protein | - | - | - | - | *Gordonia alkanivorans* strain YC-RL2 | 38% | 66.40% | 8.00E-04 | CP027114 |
| hypothetical protein | - | - | - | - | *Mycolicibacter hiberniae* JCM 13571 | 85% | 67.13% | 3.00E-28 | AP022609 |
| hypothetical protein | - | - | - | - | *Mycolicibacterium doricum* JCM 12405 | 78% | 74.63% | 3.00E-53 | AP022605 |
| hypothetical protein | - | - | - | - | *Mycolicibacter hiberniae* JCM 13571 | 23% | 72.03% | 1.00E-09 | AP022609 |
| hypothetical protein | - | - | - | - | *Mycolicibacter hiberniae* JCM 13571 | 43% | 91.45% | 3.00E-37 | AP022609 |
| hypothetical protein | - | - | - | - | MAP strain MAPK_JJ1/13 | 100% | 100.00% | 0.00E+00 | CP033909 |
| hypothetical protein | - | - | - | - | MAP strain DSM 44135 | 100% | 100.00% | 0.00E+00 | CP053068 |
| hypothetical protein | GI | GI | - | GI | MAH JP-H-1[a] | 100% | 96.19% | 0.00E+00 | AP020326 |
| hypothetical protein | - | - | - | - | MAP strain Telford[a] | 100% | 99.42% | 1.00E-78 | CP033688 |

| | | | | | | | | | |
|---|---|---|---|---|---|---|---|---|---|
| hypothetical protein | - | - | - | - | MAH strain 101115 | 100% | 90.58% | 0.00E+00 | CP040255 |
| Sulfate-binding protein | - | - | - | - | *M. colombiense* CECT 3035 | 99% | 88.60% | 0.00E+00 | CP020821 |
| putative sulfate transporter | - | - | - | - | *M. colombiense* CECT 3035 | 100% | 89.45% | 0.00E+00 | CP020821 |
| hypothetical protein | GI | - | - | GI | *Rhodococcus sp.* YL-1 | 4% | 93.94% | 0.053 | CP017299 |
| hypothetical protein | - | GI | - | GI | MAH strain OCU901s_S2_2s[a] | 100% | 99.37%% | 2.00E-155 | CP018014 |
| hypothetical protein | GI | GI | GI | GI | *M. marinum* E11 | 74% | 75.42% | 6.00E-12 | HG917972 |
| hypothetical protein | GI | GI | GI | GI | MAA strain DSM 44156 | 73% | 72.95% | 3.00E-167 | CP046507 |
| Transcriptional regulator WhiB | GI | GI | GI | GI | *M. sp.* DSM 104308 isolate 901379 | 48% | 76.00% | 2.00E-15 | LR130759 |
| hypothetical protein | GI | GI | GI | GI | *Streptomyces cinereoruber* strain ATCC 19740 | 7% | 94.12% | 0.009 | CP023693 |
| hypothetical protein | GI | GI | GI | GI | *Sorangium cellulosum* So0157-2 | 5% | 85.71% | 0.002 | CP003969 |
| hypothetical protein | GI | GI | GI | GI | *Arthrobacter sp.* AQ5-05 | 13% | 96.43% | 0.59 | CP046105 |
| hypothetical protein | GI | GI | GI | GI | *M. sp.* DSM 104308 isolate 901379 | 29% | 78.79% | 4.00E-04 | LR130759 |
| hypothetical protein | GI | GI | GI | GI | MAA strain DSM 44156 | 64% | 71.12% | 4.00E-19 | CP046507 |
| hypothetical protein | GI | GI | GI | GI | *M. mantenii* JCM 18113 | 76% | 68.65% | 4.00E-56 | AP022590 |
| Putative 8-oxo-dGTP diphosphatase 3 | GI | GI | GI | GI | *M. branderi* JCM 12687 | 83% | 75.88% | 6.00E-82 | AP022606 |
| hypothetical protein | GI | GI | GI | GI | *M. canettii* CIPT 140070010 | 99% | 74.96% | 2.00E-108 | FO203509 |
| hypothetical protein | GI | GI | GI | GI | *M. canettii* CIPT 140070010 | 71% | 75.00% | 2.00E-11 | FO203509 |
| hypothetical protein | GI | GI | GI | GI | *M. canettii* CIPT 140070010 | 81% | 70.29% | 1.00E-32 | FO203509 |
| hypothetical protein | GI | GI | GI | GI | *M. avium* strain RCAD0278 | 77% | 90.48% | 3.00E-48 | CP016396 |
| hypothetical protein | GI | GI | GI | GI | *M. mantenii* JCM 18113 | 89% | 73.43% | 9.00E-25 | AP022590 |
| hypothetical protein | GI | GI | GI | GI | MAH strain MAH11 | 60% | 75.00% | 8.00E-12 | CP035744 |
| hypothetical protein | GI | GI | GI | GI | MAA strain DSM 44156 | 52% | 79.81% | 5.00E-43 | CP046507 |
| hypothetical protein | GI | GI | GI | GI | *Burkholderia cenocepacia* strain FDAARGOS_734 | 9% | 93.75% | 0.080 | CP054817 |
| hypothetical protein | - | - | - | - | MAH strain MAH11[a] | 100% | 100% | 0.000 | CP035744 |

| | | | | | | | | | |
|---|---|---|---|---|---|---|---|---|---|
| hypothetical protein | - | - | - | - | MAH strain 101115 | 95% | 91.77% | 0.00E+00 | CP040255 |
| hypothetical protein | - | - | - | - | *M. indicus* pranii MTCC 9506 | 78% | 84.44% | 1.00E-31 | CP002275 |
| HTH-type transcriptional regulator MmpR5 | - | - | - | - | MAH strain OCU901s_S2_2s[a] | 100% | 98.78% | 0.00E+00 | CP018014 |
| hypothetical protein | - | - | - | - | MAP strain DSM 44135 | 100% | 94.72% | 0.00E+00 | CP053068 |
| hypothetical protein | GI | - | GI | - | MAA strain DSM 44156[a] | 97% | 95.69% | 0.00E+00 | CP046507 |
| hypothetical protein | - | - | - | - | MAP strain DSM 44135[a] | 99% | 99.87% | 0.00E+00 | CP053068 |
| hypothetical protein | - | - | - | - | *Mycobacteroides abscessus* strain 199 | 70% | 67.25% | 2.00E-26 | CP029076 |
| hypothetical protein | - | - | - | - | *M. canettii* CIPT 140070017 | 72% | 66.59% | 3.00E-52 | FO203510 |
| hypothetical protein | GI/Ph | Ph | - | GI | MAP strain DSM 44135 | 90% | 83.81% | 1.00E-75 | CP053068 |
| hypothetical protein | GI/Ph | Ph | Ph | GI/Ph | MAP strain DSM 44135 | 92% | 82.86% | 1.00E-62 | CP053068 |
| hypothetical protein | GI/Ph | Ph | Ph | GI/Ph | MAP strain DSM 44135 | 100% | 94.60% | 0.00E+00 | CP053068 |
| Tyrosine recombinase XerC | GI/Ph | Ph | Ph | GI/Ph | MAH strain MAC109 | 93% | 84.51% | 0.00E+00 | CP029332 |
| Tyrosine recombinase XerC | GI/Ph | Ph | Ph | GI | *M. saskatchewanense* JCM 13016 | 78% | 70.50% | 3.00E-102 | AP022573 |
| hypothetical protein | GI/Ph | Ph | Ph | GI/Ph | *Sphingomonas panacis* strain DCY99 | 23% | 79.71% | 0.002 | CP014168 |
| F420H(2)-dependent biliverdin reductase | GI | GI | GI | GI | MAH JP-H-1[a] | 98% | 98.04% | 2.00E-39 | AP020326 |
| hypothetical protein | - | - | - | - | MAA strain DSM 44135[a] | 100% | 99.33% | 2.00E-67 | CP046507 |
| Tyrosine recombinase XerC | - | - | - | - | *Mycolicibacterium sarraceniae* JCM 30395 | 99% | 78.02% | 0.00E+00 | AP022595 |
| hypothetical protein | - | - | - | - | *Burkholderia sp.* MSMB0266 | 13% | 81.54% | 0.009 | CP013417 |
| hypothetical protein | - | - | - | - | MAP strain Telford | 100% | 99.31% | 0.00E+00 | CP033688 |
| hypothetical protein | - | - | - | - | MAP strain DSM 44135 | 100% | 99.66% | 0.00E+00 | CP053068 |
| hypothetical protein | GI | - | GI | GI | MAP strain DSM 44135 | 100% | 95.46% | 0.00E+00 | CP053068 |
| hypothetical protein | GI | - | GI | GI | MAP strain DSM 44135 | 100% | 98.62% | 0.00E+00 | CP053068 |
| hypothetical protein | GI | - | GI | GI | MAP strain DSM 44135 | 100% | 98.83% | 0.00E+00 | CP053068 |
| hypothetical protein | GI | - | GI | GI | MAP strain DSM 44135 | 100% | 95.08% | 3.00E-74 | CP053068 |

| | | | | | | | | | |
|---|---|---|---|---|---|---|---|---|---|
| hypothetical protein | GI | - | GI | GI | *M. xenopi* JCM 15661T | 66% | 91.05% | 0.00E+00 | AP022314 |
| hypothetical protein | - | - | GI | - | MAH strain MAH11[a] | 99% | 100% | 3.00E-120 | CP035744 |
| hypothetical protein | - | - | GI | - | MAP strain DSM 44135 | 100% | 99.06% | 3.00E-100 | CP053068 |
| hypothetical protein | - | - | GI | - | MAA strain DSM 44135[a] | 100% | 97.38% | 2.00E-180 | CP053068 |
| hypothetical protein | - | - | - | - | MAH TH135[a] | 100% | 95.24% | 8.00E-88 | AP012555 |
| hypothetical protein | - | - | - | - | MAP strain DSM 44135 | 100% | 99.33% | 2.00E-67 | CP053068 |
| hypothetical protein | - | - | - | - | MAP strain DSM 44135 | 100% | 98.51% | 0.00E+00 | CP053068 |
| Methyl-branched lipid omega-hydroxylase | - | - | - | - | MAP strain DSM 44135 | 100% | 98.95% | 0.00E+00 | CP053068 |
| hypothetical protein | - | - | - | - | MAP strain DSM 44135 | 100% | 98.85% | 0.00E+00 | CP053068 |
| hypothetical protein | GI | GI | - | GI | MAH TH135[a] | 100% | 99.11% | 0.00E+00 | AP012555 |

[a]: The genes which had more than 95% coverage and similarity with MAH strain via BLAST analysis.

Supplementary Table 5. Geographic and host features of MAH lineages.

| Lineage | | Geographical origin | Host or niche | Notable feature of the chromosome |
|---|---|---|---|---|
| EastAsia | EA1* | Japan, Korea | Human adult, bathroom | Highly mosaic |
| | EA2* | Japan, Korea | Human adult, bathroom | Relatively few imports, inversion |
| SC1 | | USA | Little information | Little information |
| SC2/4 | SC2* | Germany, Belgium, the Netherlands, Russia, USA, Japan (pig) | Human adult and child, soil, dust, pig | Relatively few imports |
| | SC4* | | Animals, soil, dust, human adult and child | Close relative of SC2, highly mosaic |
| SC3 | | USA, Germany, Japan (pig) | Animals, water, soil, human | Highly mosaic |
| SC5 | | Japan (pig) | Pig | Close |

*: Lineage East Asia and SC2/4 was divided into two lineages respectively in the past study (21).

### References


4. T. Adachi, K. Ichikawa, T. Inagaki, M. Moriyama, T. Nakagawa, K. Ogawa, Y. Hasegawa, T. Yagi, Molecular typing and genetic characterization of *Mycobacterium avium* subsp. *hominissuis* isolates from humans and swine in Japan. J. Med. Microbiol. 65 (2016) 1289-1295.

13. K. Arikawa, T. Ichijo, S. Nakajima, Y. Nishiuchi, H. Yano, A. Tamaru, S. Yoshida, F. Maruyama, A. Ota, M. Nasu, D.A. Starkova, I. Mokrousov, O.V. Narvskaya, T. Iwamoto. Genetic relatedness of *Mycobacterium avium* subsp. *hominissuis* isolates from bathrooms of healthy volunteers, rivers, and soils in Japan with human clinical isolates from different geographical areas. Infect. Genet. Evol. 74 (2019) 103923.

14. K. Ichikawa, J. van Ingen, W.J. Koh, D. Wagner, M. Salfinger, T. Inagaki, K.I. Uchiya, T. Nakagawa, K. Ogawa, K. Yamada, T. Yagi. Genetic diversity of clinical *Mycobacterium avium* subsp. *hominissuis* and *Mycobacterium intracellulare* isolates causing pulmonary diseases recovered from different geographical regions. Infect. Genet. Evol. 36 (2015) 250-255.

16. T. Iwamoto, C. Nakajima, Y. Nishiuchi, T. Kato, S. Yoshida, N. Nakanishi, A. Tamaru, Y. Tamura, Y. Suzuki, M. Nasu. Genetic diversity of *Mycobacterium avium* subsp. *Hominissuis* strains isolated from humans, pigs, and human living environment. Infect. Genet. Evol. 12 (2012) 846-852.

19. M. Subangkit, T. Yamamoto, M. Ishida, A. Nomura, N. Yasiki, P.E. Sudaryatma, Y. Goto, T. Okabayashi. Genotyping of swine *Mycobacterium avium* subsp. *hominissuis* isolates from Kyushu, Japan. J. Vet. Med. Sci. 81 (2019) 1074-1079.

21. H. Yano, H. Suzuki, F. Maruyama, T. Iwamoto. The recombination-cold region as an epidemiological marker of recombinogenic opportunistic pathogen *Mycobacterium avium*. BMC. Genomics. 20 (2019) 752.
22. T. Komatsu, K. Ohya, A. Ota, Y. Nishiuchi, Y. Hirokazu, K. Matsuo, J.O. Odoi, S. Suganuma, K. Sawai, A. Hasebe, T. Asai, T. Yanai, H. Fukushi, T. Wada, S. Yoshida, T. Ito, K. Arikawa, M. Kawai, M. Ato, A.D. Baughn, T. Iwamoto, F. Maruyama. Genomic features of *Mycobacterium avium* subsp. *hominissuis* isolated from pigs in Japan. Gigabyte (2021) https://doi.org/10.46471/gigabyte.33.
23. T. Iwamoto, K. Arikawa, C. Nakajima, N. Nakanishi, N. Nishiuchi, Y. Yoshida, A. Tamaru, Y. Tamura, Y. Hoshino, H. Yoo, Y.K. Park, H. Saito, Y. Suzuki. Intra-subspecies sequence variability of the MACPPE12 gene in *Mycobacterium avium* subsp. *hominissuis*. Infect. Genet. Evol. 21 (2014) 479-483.
24. D.A. Starkova, T.F. Otten, I.V. Mokrousov, A.A. Viazovaia, B.I. Vishnevskiĭ, O.V. Narvskaia. Genotypic characteristics of *Mycobacterium avium* subsp. *hominissuis* strains. Genetika. 49 (2013) 1048-1054.
25. R.S. Kaas, P. Leekitcharoenphon, F.M. Aarestrup, O. Lund. Solving the problem of comparing whole bacterial genomes across different sequencing platforms. PLoS. One. 11 (2014) e104984.
26. J. Trifinopoulos, L.T. Nguyen, A. von Haeseler, B.Q. Minh. W-IQ-TREE: a fast online phylogenetic tool for maximum likelihood analysis. Nucleic. Acids. Res. 44 (2016) W232-W235.

27. G. Tonkin-Hill, J.A. Lees, S.D. Bentley, S.D.W. Frost, J. Corander. Fast hierarchical Bayesian analysis of population structure. Nucleic. Acids. Res. 47 (2019) 5539-5549.
